# Tissue-Specific Regulation of Translational Readthrough Tunes Functions of the Traffic Jam Transcription Factor

**DOI:** 10.1101/2020.12.04.411694

**Authors:** Prajwal Karki, Travis D. Carney, Cristina Maracci, Andriy S. Yatsenko, Halyna R. Shcherbata, Marina V. Rodnina

## Abstract

Translational readthrough (TR) occurs when the ribosome decodes a stop codon as a sense codon, resulting in two protein isoforms synthesized from the same mRNA. TR is pervasive in eukaryotic organisms; however, its biological significance remains unclear. In this study, we quantify the TR potential of several candidate genes in *Drosophila melanogaster* and characterize the regulation of TR in the large Maf transcription factor Traffic jam (Tj). We used CRISPR/Cas9 generated mutant flies to show that the TR-generated Tj isoform is expressed in the nuclei of a subset of neural cells of the central nervous system and is excluded from the somatic cells of gonads, which express the short Tj isoform only. Translational control of TR is critical for preservation of neuronal integrity and maintenance of reproductive health. Fine-tuning of the gene regulatory functions of transcription factors by TR provides a new potential mechanism for cell-specific regulation of gene expression.

**Highlights:**

- Tj undergoes tissue-specific TR in neural cells of the central nervous system.
- Strict control of TR is crucial for neuroprotection and maintenance of reproductive capacity.
- TR selectively fine-tunes the gene regulatory functions of the transcription factor.
- TR in Tj links transcription and translation of tissue-specific control of gene expression.

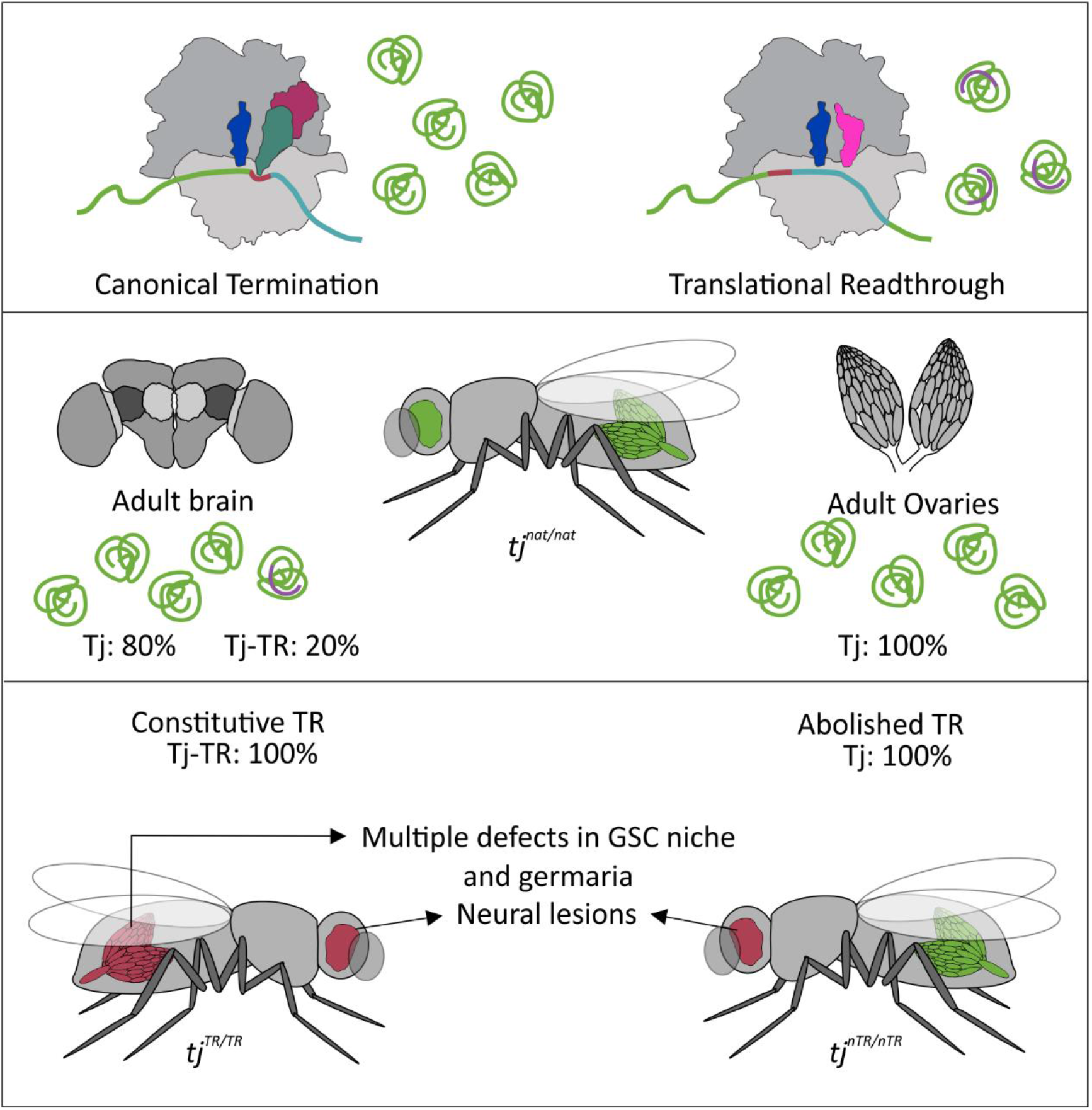

## INTRODUCTION

Eukaryotes employ several mechanisms to enlarge the coding capacity of their genomes, such as alternative splicing, alternative polyadenylation, frameshifting, and alternative initiation of translation (Kim et al., 2007; Kornblihtt et al., 2013; Tian and Manley, 2017; Touriol et al., 2003). Translational readthrough (TR) is yet another strategy to increase the diversity of the proteome by supplying C-terminally extended protein isoforms with potentially altered physiological functions. Studies over the last decade have revealed that TR can be highly pervasive in eukaryotes (Dunn et al., 2013; Jungreis et al., 2011; Loughran et al., 2014; Namy et al., 2003). The TR efficiency depends on the stop codon and its sequence context and is lowest on the UAA and highest on the UGA stop codon (Cridge et al., 2018; Howard et al., 2000; Loughran et al., 2014; Manuvakhova et al., 2000). A cytidine (C) at position +4 enhances TR (nucleotide numbering starting with the 1^st^ nucleotide of the stop codon), whereas nucleotides +4 to +9 modulate TR in a number of viral and eukaryotic genes (Beier and Grimm, 2001; Cridge et al., 2018; Loughran et al., 2014; Urban et al., 1996).

The prevalence of TR varies between organisms. It is widely employed by viruses to expand the coding potential of their small genomes (Felsenstein and Goff, 1988; Firth et al., 2011; Hofstetter et al., 1974; Pelham, 1978). Several cases of TR were described in mammalian cells, yeast, and mosquito (Dunn et al., 2013; Eswarappa et al., 2014; Jungreis et al., 2016; Namy et al., 2003; Williams et al., 2004). Analysis of the stop codon contexts of 12 *Drosophila* species and ribosome profiling studies suggested potential TR in several hundred *Drosophila* genes (Dunn et al., 2013; Jungreis et al., 2016; Jungreis et al., 2011). The majority of TR candidate genes identified in *Drosophila* have regulatory roles, suggesting that appending a functional C-terminal extension may confer conditional advantage to protein function. Computational analysis of TR protein isoforms indicates that these are mostly long, modular proteins with intrinsically disordered C-termini of low sequence complexity (Kleppe and Bornberg-Bauer, 2018; Pancsa et al., 2016). The lack of a structurally ordered C-terminus might provide conformational pliability that allows the TR extensions to perform functions without distorting the native protein.

Of several hundred candidate genes that can undergo TR in *Drosophila*, only a few have been validated experimentally. One of the early examples of a gene undergoing TR is *kelch*, which encodes a short native protein and a longer extended TR protein (Xue and Cooley, 1993). The TR efficiency of *kelch* changes through development, approaching a 1:1 ratio of the two isoforms during metamorphosis. Kelch-TR isoform is expressed in a tissue-specific manner and is particularly enriched in the imaginal discs (Robinson and Cooley, 1997), but its function is not known. Another example is the *headcase* (*hdc*) gene; the TR isoform is necessary for *Drosophila* tracheal development (Steneberg and Samakovlis, 2001). Further examples are *synapsin* (*syn*), nonsense alleles of *embryonic lethal abnormal vision* (*elav*), and *wingless* (*wg*) (Chao et al., 2003; Klagges et al., 1996; Samson et al., 1995). For the majority of the candidates, the efficiency and the biological relevance of TR are not known.

Here, we tested the TR efficiency for several candidate genes in *Drosophila*. We then selected one gene, *traffic jam* (*tj*) that shows high TR efficiency, to explore the expression and biological significance of the TR protein isoform *in vivo. tj* encodes a large Maf transcription factor that regulates gonad morphogenesis, including stem cell niche specification during ovarian development (Lai et al., 2017; Panchal et al., 2017; Wingert and DiNardo, 2015) and collective cell migration during oogenesis (Gunawan et al., 2013). Tj has extensive sequence similarity with its mammalian orthologues c-Maf and MafB, which modulate tissue-specific gene expression and cell differentiation by binding to the regulatory regions of target genes and by interacting with other transcription factors. Like all Maf factors, Tj has a leucine zipper domain, a DNA-binding basic domain, and a Maf-specific extended homology domain located at the C-terminal region of the factor (Li et al., 2003). Tj is expressed in the somatic cells of the gonads where it plays a crucial role in regulating the expression of several adhesion molecules such as Fasciclin 3 (Fas3), DE-Cadherin (DE-Cad) and Neurotactin (Nrt) (Gunawan et al., 2013; Lai et al., 2017; Li et al., 2003; Saito et al., 2009). Disruption of *tj* function causes impaired interaction between germ cells and somatic cells, which eventually leads to defective gonad development and sterility (Li et al., 2003). Apart from somatic gonadal cells, Tj is expressed in the embryonic and larval central nervous system (CNS), adult heads as well as adult fat bodies (Gelbart and Emmert, 2013; Li et al., 2003). A recent study also identified Tj as one of the transcription factors that fine-tune the molecular and morphological features of the developing neurons, contributing to remarkable glutamatergic neuronal cell-type diversity in the adult brain (Konstantinides et al., 2018).

To explore the biological significance of TR in *tj*, we utilized CRISPR/Cas9 genome editing to create mutant fly lines that exhibit different levels of TR in *tj*. Using immunohistochemistry, we show that the expression of the longer TR isoform of Tj is tissue-specific and is tightly translationally controlled. The expression of the Tj-TR isoform is important for the cells of the nervous system but is detrimental for the correct development of the gonads. Strict control of TR in a transcription factor provides a yet unidentified link between transcription and translation, which may be particularly relevant for the nervous cells that are under pressure to rapidly adapt to external stimuli and conditions.

## RESULTS

### Quantification of TR in candidate genes in *Drosophila*

Phylogenetic analysis identified more than 300 TR candidates in *Drosophila* (Jungreis et al., 2011), the majority of which have not been experimentally tested. We narrowed down the list of potential candidates to 11 genes that perform important functions during fly development (Table S1). Their functions are well characterized and associated with traceable phenotypes. Among the selected genes, *klumpfuss* (*klu*), *doublesex* (*dsx*), *traffic jam* (*tj*), *seven up* (*svp*), *chronologically inappropriate morphogenesis* (*chinmo*), *fruitless* (*fru*), and *broad* (*br*) encode transcription factors or transcriptional regulators; *atypical protein kinase C* (*aPKC*), and *discs large 1* (*dlg1*) encode protein kinases involved in cell signaling; *wishful thinking* (*wit*) encodes a signaling receptor; and *kinesin heavy chain-73* (*khc-73*) encodes a motor protein that regulates cell polarity. The lengths of the expected TR extensions range from 11 to 236 amino acids (Table S1). With the exception of *wit*, the respective mRNAs do not have a propensity to form secondary structures in their 3’ UTR as predicted by RNAfold.

To validate the TR of the candidate genes, we constructed dual-luciferase reporters for the 11 candidate genes and tested their expression in S2 cell lines (Figure 1A). We generated a psiCHECK^™^-2 based dual-luciferase vector where the *Renilla* and *Firefly* luciferase genes are fused into a single ORF, separated by the test cassette for each gene and a self-cleaving P2A sequence. For each gene construct, we generated a positive control by mutating the native stop codons to the UUC sense codon, resulting in a constitutive Firefly synthesis. Additionally, we mutated the extended tetranucleotide stop codon sequence to UAA-A to obtain control constructs with a highly efficient translation termination context. *αTubulin 84B* (*αTub84B*), which is not a TR candidate, was used as a negative control. The constructs were introduced into S2 cells, and TR efficiency was calculated from the activity of Renilla and Firefly reporters. TR efficiency of the control *αTub84B* sequence is 0.35% (Figure 1B), similar to basal TR frequency of 0.02-1.4% reported for yeast and mammalian cell lines (Bonetti et al., 1995; Fearon et al., 1994; Firth et al., 2011; Keeling et al., 2004; Namy et al., 2002; Napthine et al., 2012). Mutating the native tetranucleotide termination signal of *αTub84B* from UAA-G to UGA-C does not increase TR values, indicating that the 105 bp *αTub84B* test cassette represents a robust sequence with efficient termination independent of the immediate stop codon context.

**Figure 1.**
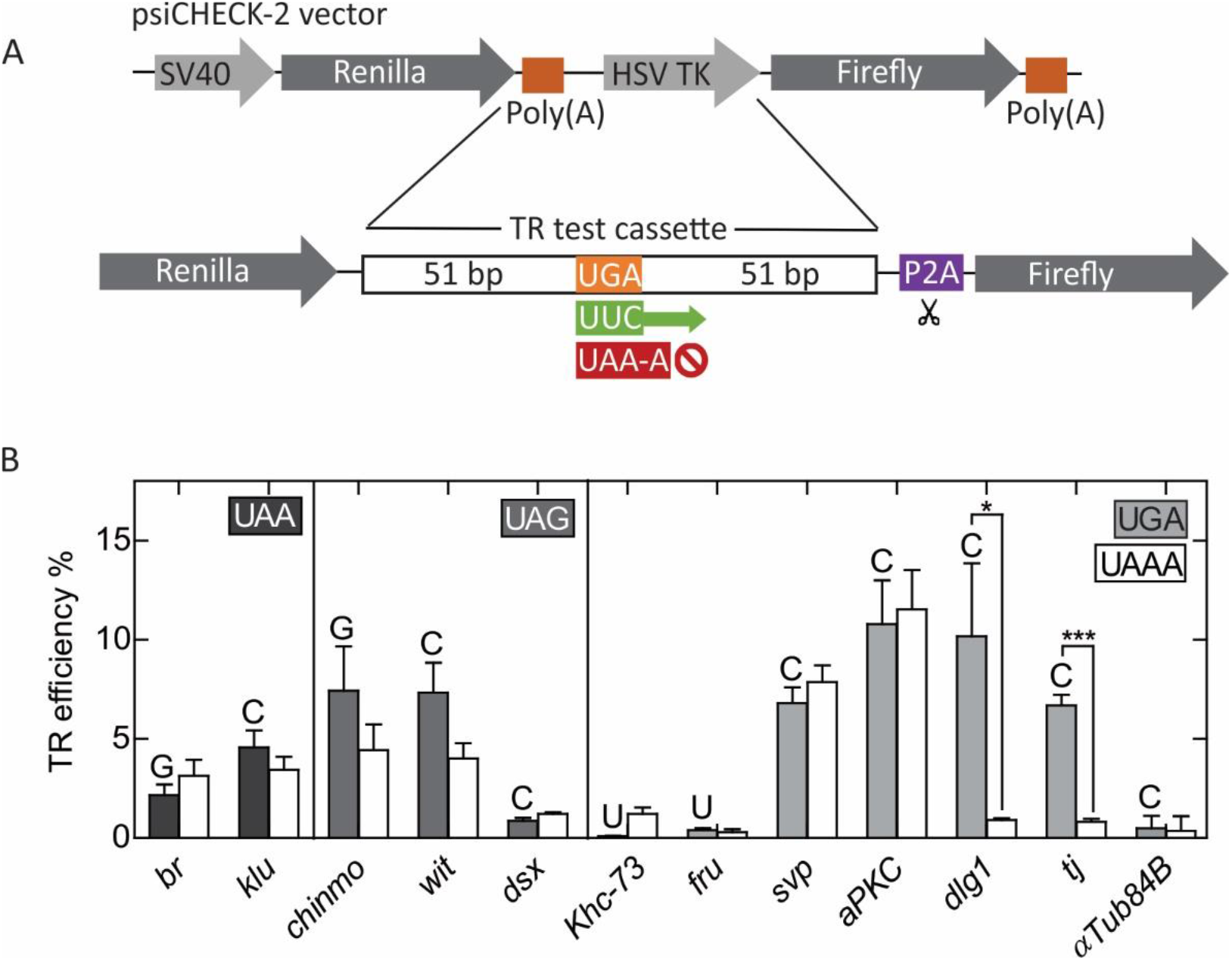
Quantification of TR in putative candidate genes using dual luciferase reporter assay in S2 cells. (A) Construct design for dual luciferase reporter vectors. (B) TR efficiencies of putative candidate genes with UAA, UAG, and UGA stop codons determined by dual luciferase assay in S2 cells. The +4 nucleotide for each gene is indicated by the letter above each bar. White bars represent TR efficiencies for corresponding genes upon mutating the native tetranucleotide termination signal to UAA-A. *αTub84B* represents a negative control. The error bars indicate the SD from three biological replicates. p-values are calculated using two-tailed unpaired Student’s t-test from datasets generated from three biological replicates, each with three technical replicates. (*p<0.05, ***p<0.0005). See also Table S1.

Among the candidate genes selected, *dsx*, *khc73*, and *fru* show basal TR levels, *i.e*. they do not undergo TR at the conditions tested (Figure 1B). The TR efficiency of *br* and *klu*, which harbor a UAA stop codon followed by G or C, is independent of the nucleotide at +4 position, suggesting that the moderate TR may be induced by elements beyond the stop codon. For genes containing a UAG stop codon, *chinmo* and *wit*, the TR efficiency is reduced by about a half by mutating nucleotides +3 and +4, suggesting that TR depends not only on the extended stop codon, but also on an external signal(s). The four genes with the UGA-C stop signal, *svp*, *aPKC, dlg1* and *tj*, exhibit the highest TR efficiencies, ranging from about 7% to 11%. Mutating tetranucleotide signal to UAA-A in *dlg1* and *tj* abolishes TR efficiency, indicating that the immediate nucleotide context is the only requirement to drive TR in these genes. TR in *svp* and *aPKC* is unaffected upon mutating the tetranucleotide termination signal to UAA-A.

### Tissue-specific TR in *tj* during embryogenesis and in adult *Drosophila*

To study the physiological relevance of TR in *Drosophila*, we have chosen *tj* as a model TR gene. The leaky UGA-C tetranucleotide is sufficient to induce TR in *tj* (Figure 1B). Because the gene encoding *tj* lacks introns, genetic manipulation of the gene is relatively simple and avoids the complications of working with multiple splice isoforms. The ORF of *tj* is 509 codons-long; the TR extension would append additional 44 amino acids, generating a larger protein that we call the Tj-TR isoform.

To study TR in *Drosophila*, we created three mutant fly lines that harbor mutations at the termination sequence of *tj* using CRISPR/Cas9-based genome editing (Figure S1). The mutants were designed to code for a Flag epitope tag downstream of the TR extension (Figure 2A). The first mutation, *tj^nat^*, does not alter the primary stop signal that terminates the *tj* ORF; TR in this mutant is expected to occur at the same frequency as in the native *tj*. The *tj^TR^* mutation replaces the primary *tj* stop codon with a UUC sense codon, such that the mutant flies undergo constitutive TR and produce only the Flag-tagged TR isoform. Finally, the *tj^nTR^* mutation introduces multiple stop codons after the primary *tj* stop codon, which leads to complete abolition of TR. Flies homozygous for each of the three genomic *tj*-TR mutations exhibit no gross defects in growth and viability.

**Figure 2.**
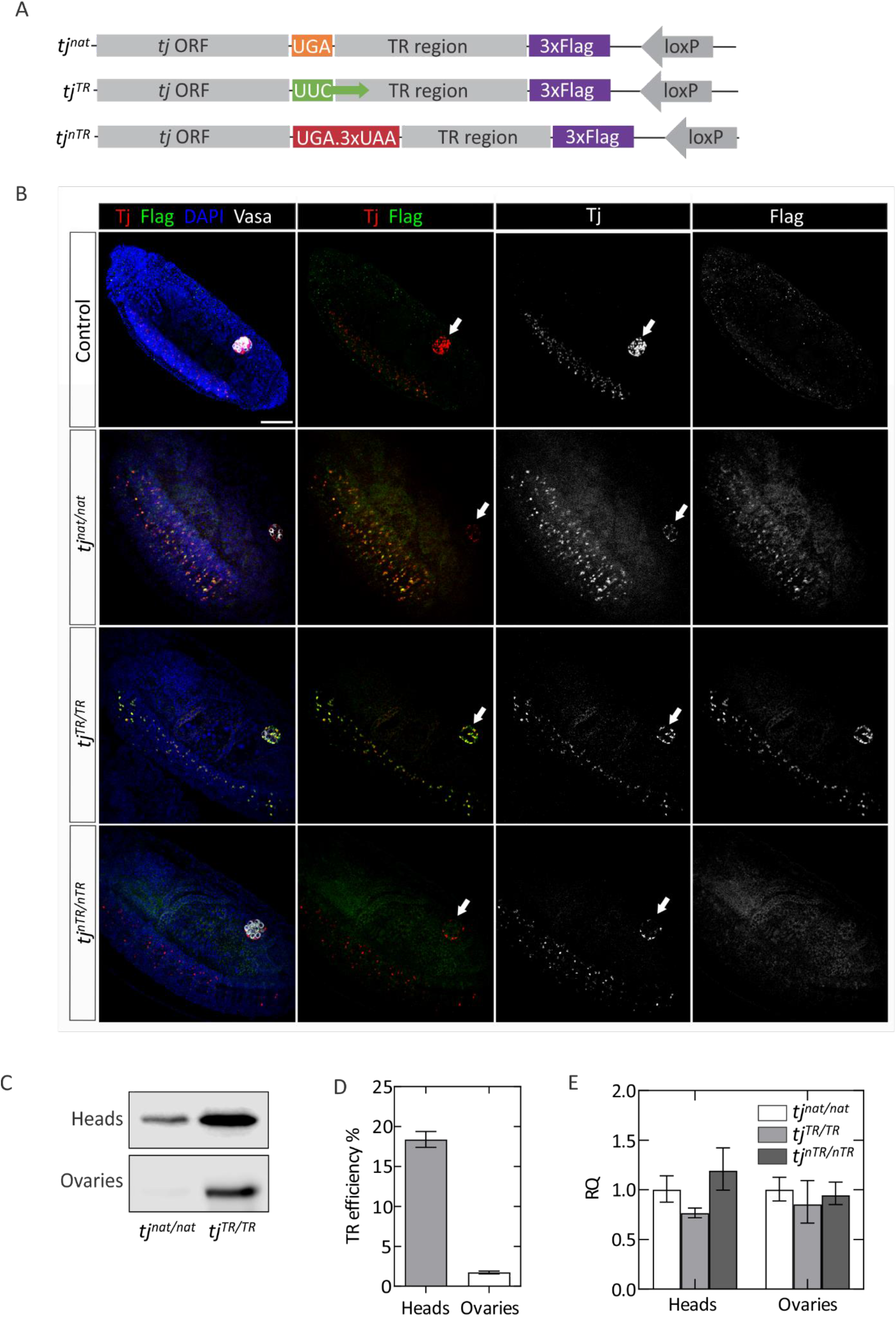
Tissue-specific expression of the Tj-TR isoform in Drosophila. (A) Sequence map of chromosomal modifications introduced in the *tj* locus by CRISPR/Cas9 genome editing. (B) Tissue-specific regulation of TR in *tj* during embryogenesis. Stage 15-17 embryos are stained with the nuclear stain DAPI (blue), anti-Tj (red), and anti-Flag (green) antibodies, as well as anti-Vasa (greyscale) to mark the gonads. Individual channels for anti-Tj (grey) and anti-Flag (grey) and the overlapping channel are shown in separate columns. In control (first row) and in *tj^nat/nat^* (second row), embryos express Tj in neural cells of embryonic central nervous system (CNS, visible as the row of Tj-positive cells near the left of each panel) as well as in somatic gonadal precursor cells (SGPs) of embryonic gonads (arrowheads). The Flag-tagged Tj-TR isoform is selectively expressed in the embryonic VNC and excluded from gonads. *tj^TR/TR^* embryos (third row) exhibit constitutive expression of the Tj-TR isoform in both tissues while *tj^nTR/nTR^* embryos (fourth row) do not express the Tj-TR isoform in any embryonic tissues. Scale bar: 50 μm. (C) Western blot with anti-Flag antibodies showing the relative abundance of Flag-tagged Tj-TR isoform in adult tissues from *tj^nat/nat^* and *tj^TR/TR^* mutants. (D) TR efficiency estimated in adult heads and ovaries using quantitative western blot using total protein normalization (STAR Methods). (E) RT-qPCR analysis of *tj* transcripts using cDNA prepared from adult heads of *tj^nat/nat^* and *tj^TR/TR^* mutants. Error bars represent the upper and lower limit of RQ defined by the standard deviation of ΔΔC_T_ from three biological replicates. The data were normalized against average ΔC_T_ of the housekeeping gene *αTub84B*. Two-tailed unpaired Student’s t-test performed in both the datasets indicate that the differences are statistically non-significant. See also Figures S1, S2 and S3.

To detect the expression of Tj and Tj-TR isoforms during embryonic development, we stained stage 15-17 embryos with antibodies specific to Tj and Flag to visualize Tj and Tj-TR isoforms, respectively. In controls and in all *tj-TR* mutant embryos, Tj is expressed in a subset of neural cells in the ventral nerve cord (VNC) and brain as well as in the somatic gonadal precursors (SGPs) of the embryonic gonad (Figures 2B and S3A). The Tj-TR isoform is expressed in the VNC in *tj^nat/nat^* embryos (Figure 2B). Interestingly, we could not detect the Tj-TR isoform in the embryonic gonads of the same mutants. *tj^TR/TR^* mutants constitutively express the Tj-TR isoform in the VNC as well as gonads, while *tj^nTR/nTR^* mutants only express the native Tj. Selective expression of the Tj-TR isoform in the nervous system of *tj^nat/nat^* mutants suggests that TR in *tj* is regulated in a tissue-specific manner during embryogenesis. Furthermore, the signals from Tj and Tj-TR isoforms overlap perfectly with the DAPI stain in the embryonic SGPs (Figure S3A), indicating that the TR extension does not affect the nuclear localization of Tj.

To estimate the TR efficiency in adult tissues, we utilized quantitative western blot and probed the abundance of Flag-tagged Tj-TR isoform in tissue lysates from *tj^nat/nat^* and *tj^TR/TR^* adult flies using anti-Flag antibodies. We estimated a TR efficiency of ~20% in adult heads, whereas adult ovaries exhibited only basal levels of TR (Figures 2C, D). The potential differences in *tj* transcript levels between the mutants were ruled out by RT-qPCR experiments (Figure 2E). Given that the Tj staining is very similar in both *tj^TR/TR^* and *tj^nTR/nTR^* mutant lines (Figures S2 and S3), the TR extension does not seem to trigger selective degradation of the protein. Thus, TR is regulated on a translational level and is not caused by tissue-specific differences in mRNA levels or protein stability or by altered cellular localization of the two Tj isoforms.

### Effect of TR induction in adult brains

To test whether the nervous tissue-specific expression of Tj-TR isoform persists until adulthood, we inspected adult brains from all three *tj*-TR mutants. The brain is composed of two bilateral hemispheres, each of which is divided into a medial compartment termed the central brain (CB) and a lateral optic lobe (OL), which receives primary visual input from photoreceptor neurons. Marking *tj*-expressing cells with membrane-bound RFP (*tj>mRFP*) allows the visualization of neuronal processes and demonstrates that the brain contains many Tj-positive neurons, especially dispersed throughout the OL; in addition, several prominent clusters of Tj-expressing cells exist, particularly in the junction of the lobula and the CB and in some neurons of the pars intercerebralis (PI) (Figure S2A). Next, we examined brains from control as well as all three *tj*-TR mutants (Fig S2B-E). The pattern and number of Tj-expressing cells does not appear to be different in any of these genotypes. Furthermore, the cells expressing Tj in the brains of *tj^nat/nat^* flies also express the Tj-TR isoform, indicating that the nervous tissue-specific regulation of TR in *tj* is maintained in adults (Figure S2C). As expected, in *tj^TR/TR^* brains, Flag expression exhibits complete overlap with Tj (Fig S2D), while Flag expression is absent from *tj^nTR/nTR^* brains (Fig S2E).

Because Tj-TR is expressed in the nervous system (Figures 2B and S2), we analyzed the effect of differential TR in adult brains. For all mutants, we analyzed brains from young (3-4 day old) as well as aged (one month old) adults to examine the possibility of age-related defects. We also included brains from *tj* hypomorphic flies (*tj^hypo^*) to compare with the *tj*-TR mutants. The hypomorphic flies bore the strong hypomorphic *tj^PL3^* allele in trans to *tj-Gal4*, which itself is reported to be a weak hypomorphic allele (Panchal et al., 2017). We stained brains with an antibody against the cell adhesion molecule DE-Cadherin (Cad), whose ubiquitous expression allows visualization of the neuropils of the brain. In addition, we prepared brain sections stained with hematoxylin and eosin (H&E) for histological evaluation. Using these methods, we found evidence of neural lesions in *tj^hypo^* brains as well as in both *tj^TR/TR^* and *tj^nTR/nTR^* brains (Figure 3; Table S3). In immunofluorescence images, these lesions are visible either as regions in which Cad staining is cleared, indicating the absence of neural cells or projections, or as regions of “sponge-like” tissue containing sporadic, small lesions (Figure 3A). In H&E sections, lesions are visible as stain-free clearings (Figure 3D). The incidence of these lesions is significantly greater in brains of *tj^TR/TR^* and *tj^nTR/nTR^* flies than in *tj^nat^* (Figures 3B and 3C). As the majority of Tj-expressing cells reside in the OL, we sought to determine whether the incidence of lesions is greater in the OL than in the CB. We quantified brain lesions independently in these compartments and found that their incidence is very similar in both (Figures 3B and 3C; Table S3). This is consistent with the innervation of the CB by OL neurons. To determine if the lesions worsen with age, we also performed these analyses on aged, one-month old brains. We found that the incidence of lesions does not increase with age, indicating that they arise developmentally and are likely not degenerative (Table S3). These results show that in the CNS, regulation of Tj and Tj-TR expression is crucial for neuroprotection.

**Figure 3.**
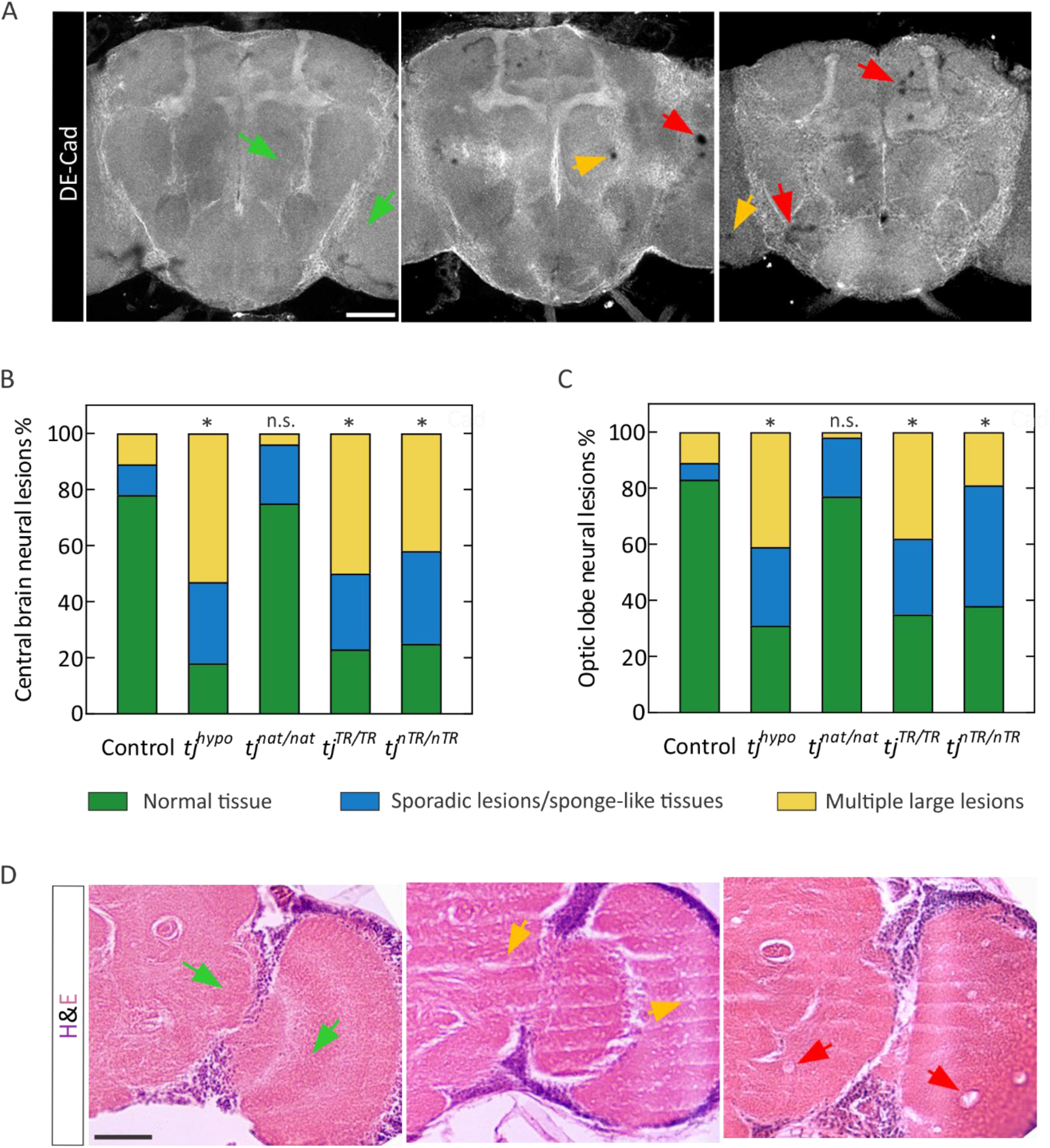
Neural lesions in adult brains caused by perturbation of proper regulation of *tj*-TR. (A) Examples of neural lesion phenotypes in young (3-4 day old) adult brains stained with DE-Cadherin (Cad). Cad staining normally appears homogeneous (e.g., green arrowheads) and is disrupted in instances of sporadic lesions and sponge-like neural tissues (yellow arrowheads) or occasionally large lesions (red arrowheads). Focal planes were chosen to illustrate lesions, which occur throughout *tj^hypo^, tj^TR/TR^*, and *tj^nTR/nTR^* brains. Scale bar: 100 μm. (B) Quantification of phenotypes in central brains. (C) Quantification of phenotypes in optic lobes. Two-way tables and Chi-square tests were used to determine any significant differences in the distribution of phenotypes between the mutants (*p <.05; n.s., not significantly different). (D) Hematoxylin and eosin (H&E)-stained brain sections chosen to illustrate wild-type appearance (green arrowheads), sporadic lesions (yellow arrowheads), and large lesions (red arrowheads). Scale bar: 100 μm. See also Figure S2 and Table S3.

### Effect of TR induction in ovaries

As *tj* plays an important role in gonad development, we specifically examined how TR affects Tj function in adult ovaries. Each ovary in *Drosophila* is composed of about nineteen parallel egg production units called the ovarioles. At the anterior tip of each ovariole is a structure called the germarium that houses the germline stem cells (GSCs) and the GSC niche (Figure 4A). The GSC niche comprises an anterior stack of disc-shaped terminal filament cells (TFCs) and a cluster of 6-7 cap cells (CpCs) that make direct contact with GSCs (Yatsenko and Shcherbata, 2018). GSCs divide asymmetrically to generate cystoblasts that further divide to yield a 16-cell germline cluster. These clusters become enveloped by somatic follicle cells to form cysts that bud off from the posterior end of the germarium. Germline cells can be identified based on expression of Vasa, an RNA-binding protein. TFCs and CpCs express the transcription factor Engrailed (En). Tj is expressed in CpCs but not in the TFCs (Li et al., 2003; Panchal et al., 2017). Tj is also expressed in some other somatic cells, including the follicle cells that surround the germline cysts, as well as escort cells (ECs), which are interspersed with and comprise a differentiation niche for germline cells in the germarium (Figure S3) (Li et al., 2003; Panchal et al., 2017). The cell adhesion molecule Fas3 is expressed in the follicle cells in early egg chambers as well (Bai and Montell, 2002).

**Figure 4.**
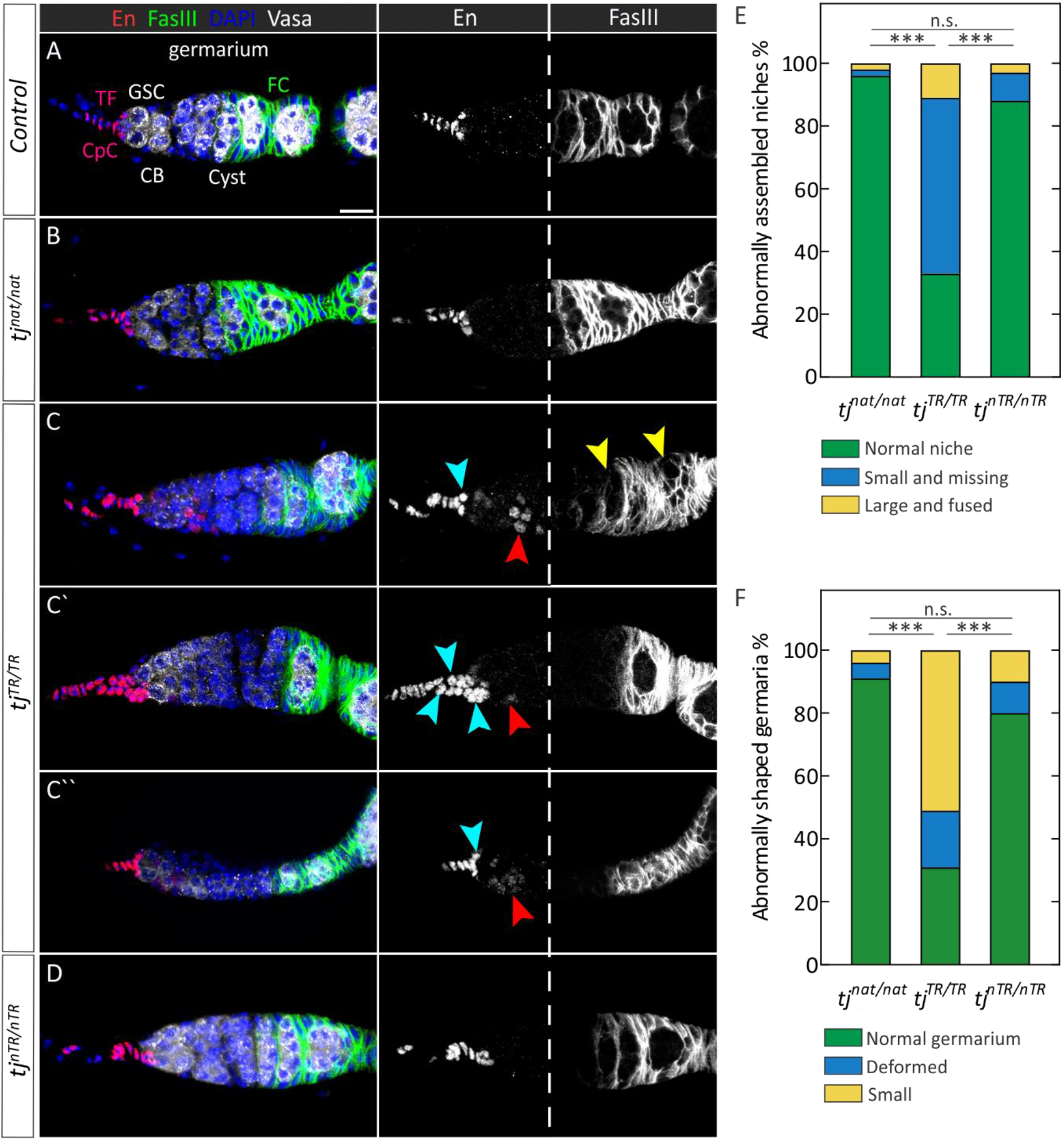
Defects in adult germaria due to constitutive TR in Tj. (A) Control germarium. Engrailed (En) is expressed in terminal filament cells (TFCs) and CpCs. Fas3 is present in follicle cells, which are present in the posterior half of the germarium (right panel from the white dashed line) and in the follicular epithelia surrounding the early-stage egg chambers. Shown are some of the relevant cell types (CB, cystoblast; CpC, cap cell; FC, follicle cell; GSC, germline stem cell; TF, terminal filament). Scale bar: 10 μm. (B) Wild-type-like morphology of *tj^nat/nat^* germaria. (C-C’’) Diverse phenotypes observed in *tj^TR/TR^* niches and germaria. Some niches appear small (blue arrowheads, C and C’’), some are enlarged, appearing to be composed of multiple niches that have fused (multiple blue arrowheads, C’), while some are indiscernible or absent. Similarly, a majority of the germaria are small or deformed (C’’). Ectopic expression of En is observed in a subset of escort cells in all *tj^TR/TR^* germaria (red arrowheads). In some germaria, the Fas3-positive follicle cells fail to encapsulate the germline cells (yellow arrowheads). (D) Organization of *tj^nTR/nTR^* germaria. The GSC niche and germaria of *tj^nTR/nTR^* ovaries appear normal and are indistinguishable from control or *tj^nat/nat^*. (E) Quantification of niche phenotypes described above and observed in (D-F). (F) Quantification of defects in germaria observed in (B-D). Two-way tables and Chi-square tests were used to determine any significant differences in the distribution of phenotypes between the mutants (***, p < 10^-4^; n.s., not significantly different). See also Figures S3, S4, and Table S2.

Using En as a marker for TFCs and CpCs, it is possible to distinguish between them based on the disc shape and stacked organization of the TFCs and the clustered, round, germline-adjacent nature of CpCs. Thus, we used En staining and determined that the GSC niches appear indistinguishable from control in the *tj^nat/nat^* and *tj^nTR/nTR^* flies (Figure 4), a result consistent with our observation that TR is absent in gonads. In contrast, constitutive TR in *tj* produces diverse defects (Figures 4C–4C”). Whereas *tj^nat/nat^* and *tj^nTR/nTR^* germaria nearly always have normal terminal filaments and a cluster of about six CpCs (Figures 4A and 4D), *tj^TR/TR^* niches are often small or absent (Figures 4C and 4C”, blue arrowheads). In addition, a small number of mutant niches appear enlarged, perhaps resulting from two or more adjacent niches that have fused (Figures 4C’, blue arrowheads; quantified in Figure 4E and Table S2).

Germaria in the *tj^TR/TR^* mutants are frequently small or exhibit deformities (Figures 4C” and Figure S4; quantified in Figure 4F and Table S2); these phenotypes are similar to those in *tj^hypo^* ovarioles (Figures S4B and S4C). While in *tj^nat/nat^* and *tj^nTR/nTR^*, En was restricted to the TFCs and CpCs, we found that 100% of *tj^TR/TR^* germaria exhibit ectopic En expression in escort cells, albeit more weakly than in the GSC niche cells (Figures 4C–4C”, red arrowheads). These results are in agreement with previous observations (Li et al., 2019), in which somatic knockdown of *tj* resulted in derepression of En in anterior ECs, which form the differentiation niche regulating the efficiency of GSC progeny differentiation (Konig and Shcherbata, 2015). Since En is an activator of *dpp* expression (Eliazer et al., 2014), its presence in ECs can disrupt the differentiation niche and lead to a failure of the germline to develop properly. Thus, our data suggest that the Tj-TR protein behaves as a hypomorphic mutant in the somatic cells of the germarium.

In the posterior part of the germarium, germline becomes encapsulated by follicle cells, which are derived from two follicle stem cells (FSCs), located about halfway along the A-P axis of the germarium. Fas3 is expressed in early follicle cells and can be used to visualize the morphology and organization of the follicular epithelium (Figure 4A). Using this marker, we frequently saw *tj^TR/TR^* germaria in which follicle cells fail to surround the germline (Figures 4C, yellow arrowheads), whereas in control, *tj^nat/nat^*, and *tj^nTR/nTR^* germaria, germline cysts are consistently encapsulated by Fas3-positive cells (Figures 4A, 4B, and 4D). Germline left unprotected by the follicular epithelium is known be susceptible to cell death; indeed, *tj^TR/TR^* ovarioles have a strong increase in the number of dying egg chambers (Figure S4D and Table S2).

### TR modulates selective gene regulatory properties of Tj

Because the Tj-TR isoform is specific to neural cells of CNS, we studied how induction or disruption of TR affects the expression of the known target genes regulated by Tj. We performed RT-qPCR with adult head tissues from all three *tj*-TR mutants. The expression levels of the *VGlut, Rh6, melt* and *wts*, which are regulated by Tj (Jukam et al., 2013; Konstantinides et al., 2018; Li et al., 2003), did not change significantly (Figure S5). High-throughput RNA sequencing (RNA-seq) on adult brain samples from the three *tj*-TR mutants identified genes deregulated in *tj^TR/TR^* or *tj^nTR/nTR^* brain samples. Included are genes that act in metabolic processes, stress responses, signaling pathways, and mitochondria (Table S4, Figure S5). We then validated the selective deregulation of several candidate target genes identified by RNA-seq using RT-qPCR analysis. Genes that exhibit marked upregulation upon TR induction in *tj* include *Inwardly rectifying potassium channel 3 (Irk3), sandman (sand), neither activation nor afterpotential D (ninaD)*, and *Odorant-binding protein 99a (Obp99a)* (Figure 5A). These genes represent potential transcriptional targets that might be positively regulated by the Tj-TR isoform. This suggests that the Tj-TR isoform affects CNS functions related to perception of external stimuli (such as light and olfactory molecules) and homeostatic cellular response to such stimuli. The observation that the expression levels of these genes in *tj^nTR/nTR^* mutants were comparable to *tj^nat/nat^* suggests that the effects are due to the overexpression of the Tj-TR isoform in *tj^TR/TR^* flies, whereas a moderate Tj-TR expression in controlled lab conditions do not upregulate these genes. Although, it is possible that an effect could also be masked by using the whole heads instead of isolated brains.

**Figure 5.**
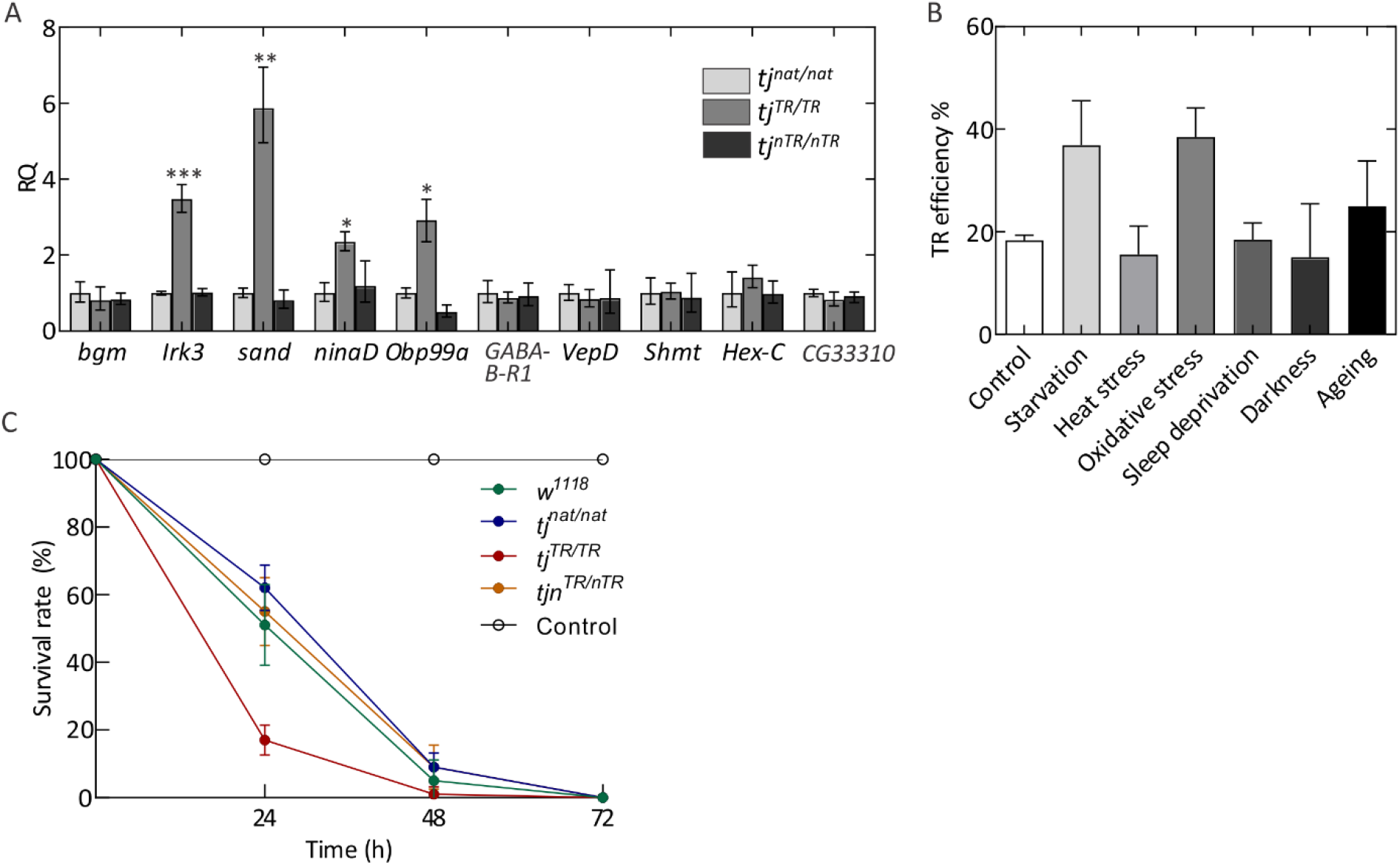
Effect of TR in tj on the transcriptome profile in adult heads. (A) RT-qPCR analysis of genes deregulated in *tj*-TR mutants using cDNA prepared from adult heads. Error bars represent the upper and lower limits of RQ values defined by the standard deviation of ΔΔC_T_. The data were normalized against average ΔC_T_ of the housekeeping gene *αTub84B*. p-values were calculated using two-tailed unpaired Student’s t-test with Welch’s correction of standard deviation from ΔΔC_T_ values of *tj^nat/nat^* and *tj^TR/TR^* samples, obtained from three biological replicates (*p<0.05, **p<0.005, ***p<0.0005). (B) Effect of different stress conditions on TR efficiencies in Tj. Head lysates from stressed *tj^nat/nat^* and *tj^TR/TR^* mutant flies were used to perform western blot. Total protein normalization was used for determining TR efficiencies. Error bars represent SD from three biological replicates. (C) Survival rate of wild-type (*w^1118^*), *tj^nat/nat^, tj^TR/TR^* and *tj^nTR/nTR^* mutants upon stress induction via paraquat ingestion (20 mM paraquat in 3% sucrose). Control group represents *w^1118^* flies that were not exposed to paraquat. Error bars represent SD from five biological replicates. See also Figure S5 and Table S4.

Because several genes upregulated in *tj^TR/TR^* mutants are largely involved in homeostatic functions, we tested whether the TR efficiency is affected by different stress conditions. We subjected *tj^nat/nat^* and *tj^TR/TR^* mutants to several stressors such as starvation, heat stress, oxidative stress, sleep deprivation and constant exposure to darkness and measured TR efficiency in adult heads using quantitative western blot. We also aged the adult flies for two weeks to test whether TR efficiency changes with chronological ageing. While heat stress, sleep deprivation, and darkness do not have an effect on TR in Tj, starvation and oxidative stress enhance the TR efficiency to approximately 40%; ageing has only a modest effect (Figure 5B). Thus, the expression level of Tj-TR can be fine-tuned at certain stress conditions.

In order to understand the physiological relevance of these observations, we tested the effect of stress induction on the survival rate of 4-5 days old wild-type and mutant flies (Figure 5C). We induced oxidative stress via paraquat ingestion and recorded the viability of flies over three days. Interestingly, the survival rates of the *tj^nTR/nTR^* mutants was comparable to that of *tj^nat/nat^* and wild-type flies, while the *tj^TR/TR^* mutants exhibited reduced survival rates. Although we did not observe any susceptibility in *tj^nTR/nTR^* flies, we cannot rule out the possibility that TR in Tj might aid in internal cellular responses that do not manifest in organismal survivability. On the other hand, the importance of a proper regulation of TR is further corroborated by the observation that *tj^TR/TR^* mutants have decreased viability when subject to acute oxidative stress.

### Mechanism of tissue-specific TR regulation

Tissue-specific differences in TR can be attributed to factors that can potentially upregulate global TR at leaky stop codons or *trans* factors that drive gene-specific TR by binding to target mRNA elements outside the stop codon (Eswarappa et al., 2014). In particular, differences in the relative abundance, activity and modifications of eukaryotic release factors (eRF1 and eRF3) and tRNAs that can recognize stop codons can play a role in modulating tissue-specific TR (Beznoskova et al., 2016; Blanchet et al., 2014; Chauvin et al., 2005). As our luciferase data suggest that TR in *tj* is solely determined by the tetranucleotide termination signal, and is independent of extended mRNA element (Figure 1B), the most likely tissue-specific regulators are those interacting with the stop codon directly.

To test this hypothesis, we first quantified the relative levels of eukaryotic release factors, eRF1 and eRF3. *eRF1* in *Drosophila* has 8 annotated transcript isoforms with at least 3 different predicted protein isoforms (Figure 6A). These protein isoforms have a conserved core but possess differentially spliced C-termini. Isoforms A, B, C, E, F and G possess the same C-terminus and give rise to a 438 aa long protein product (henceforth, referred to as eRF1A). *eRF1H* and *eRF1I* are unique isoforms that are 437 aa and 447 aa in length, respectively. *eRF3* has 4 annotated transcript isoforms (Figure 6B): isoforms A, B and D give rise to full length eRF3 (619 amino acids) and have a high degree of sequence similarity (henceforth referred to as *eRF3A*), whereas isoform C possesses an N-terminal truncation of 124 aa. We performed RT-qPCR based relative quantification of each of these factors and their isoforms by using isoform-specific primer pairs. *eRF1H* isoform was significantly overexpressed in heads (~130 folds) compared to the ovaries, while the relative abundance of *eRF1A* was comparable (Figure 6C). We note that the *eRF1* transcript pool in ovaries comprised almost entirely of *eRF1A*, whereas in the heads *eRF1A* and *eRF1H* transcripts constituted 66.8% and 33.1%, respectively, of the total eRF1 transcript pool. Additionally, *eRF1I* isoform was 10 times more abundant in heads compared to the ovaries, albeit its expression is significantly lower compared to other isoforms. In case of eRF3, the fulllength isoform, *eRF3A* was relatively less abundant in heads, and the N-terminally truncated *eRF3C* isoform was 3 times more abundant in heads compared to the ovaries (Figure 6D).

**Figure 6.**
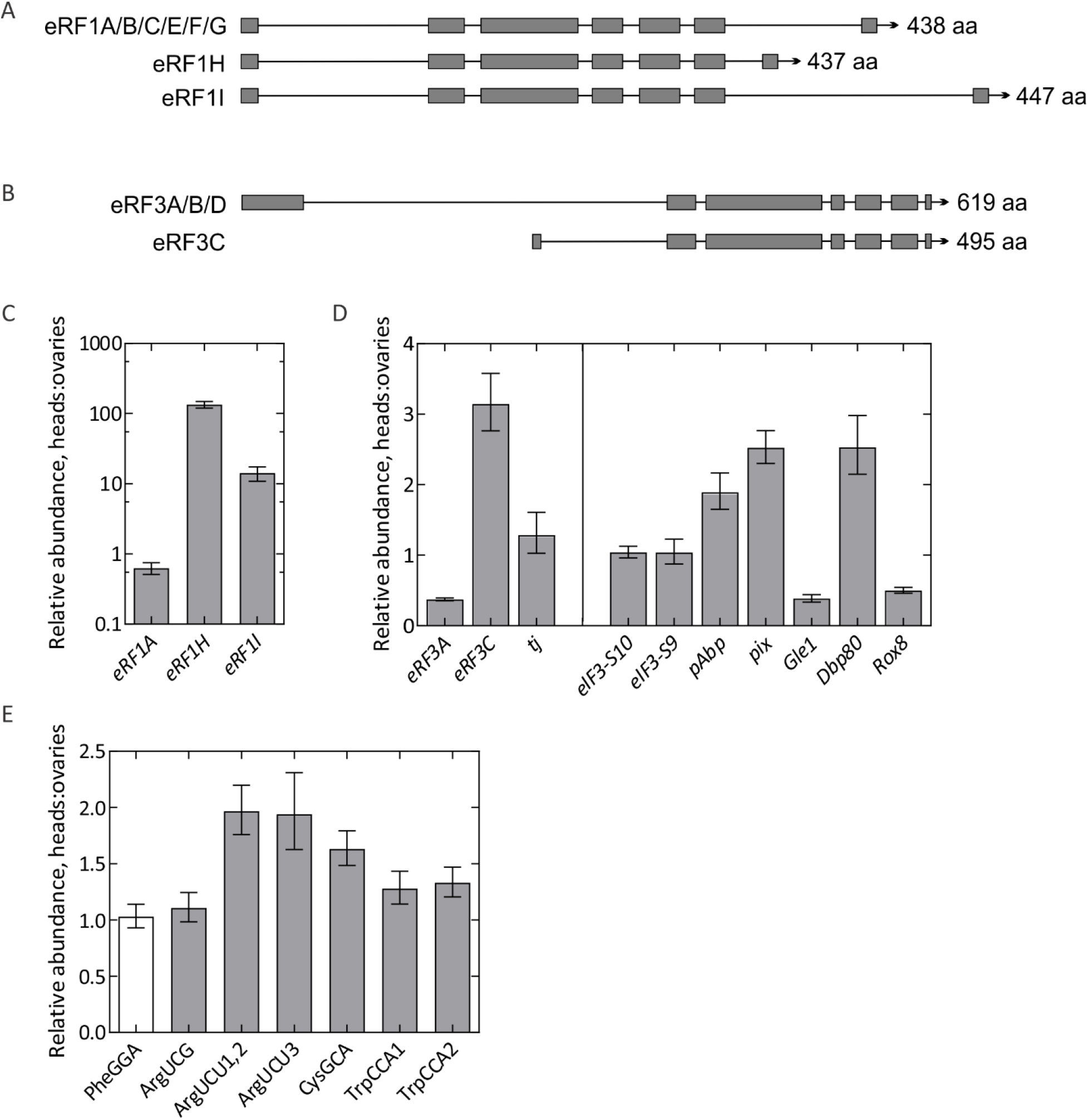
Tissue-specific differences in factors directly involved in leaky termination. Genome map depicting the ORFs of different isoforms of eRF1 (A) and eRF3 (B). (C) RT-qPCR analysis of eRF1 isoforms. Isoforms A, B, C, E, F and G are collectively depicted as *eRF1A* for simplicity. (D) RT-qPCR analysis of full-length *eRF3* (*A,B,D*), depicted as *eRF3A*; N-terminally truncated isoform *eRF3C* and *tj* (left panel); and other accessory factors implicated in termination fidelity (right panel): *eIF3* subunits *S10* and *S9, pAbp, pix, Gle1, Dbp80* and *Rox8* transcripts using cDNA prepared from adult fly heads and ovaries. C_T_ values for each transcript were normalized against the respective C_T_ values for *αTub84B*. (E) RT-qPCR analysis of ArgUCG, ArgUCU, CysGCA and TrpCCA isoacceptor tRNAs that are near-cognate to UGA stop codon. PheGGA serves as a control tRNA which is non-cognate to UGA. For cases where individual isodecoders are quantified using separate primer pairs, the isodecoder identity is indicated by the number at the end of the tRNA. C_T_ values for each transcript were normalized against the respective C_T_ values for 18S rRNA. The ΔC_T_ values obtained from each test transcript were then compared between the tissues to derive ΔΔC_T_. Error bars represent the upper and lower limit of RQ defined by the standard deviation of ΔΔC_T_ from three biological replicates.

Additionally, we also tested the relative abundance of *Drosophila* orthologues of several factors that are directly or indirectly implicated in eukaryotic translation termination fidelity and ribosome recycling. eIF3 has been reported to increase TR by interfering with eRF1 decoding of stop codon at the third/wobble position (Beznoskova et al., 2015). Rli1/ABCE1 facilitates recruitment of eRFs to the ribosome and promotes termination (Beissel et al., 2019; Shoemaker and Green, 2011). Dbp5/DDX19 stabilizes termination complex and prevents premature dissociation of eRFs (Beissel et al., 2019; Mikhailova et al., 2017), while Gle1 functions together with Dbp5 to regulate termination (Bolger et al., 2008). Pub1 and poly(A) binding protein (PABP) are known to interact with eRFs and stimulate termination efficiency (Ivanov et al., 2016; Urakov et al., 2017). Upon quantification, we did not find any difference in the relative abundance of *eIF3* (*S10* and *S9*, orthologous to *eIF3a* and *eIF3b*) between heads and ovaries. However, *pAbp* (*PABP*), *pix* (*Rli1/ABCE1*) and *Dbp80* (*Dbp5/DDX19*) were found to be 2-3 times more abundant in heads while *Gle1* and *Rox8* (*Pub1*) were relatively less abundant in heads (Figure 6E).

Another potential mechanism for tissue-specific TR regulation is the relative abundance of near-cognate tRNAs (nc-tRNAs) that can read stop codons due to wobble interactions with the nucleotide at the 1^st^ or 3^rd^ codon position. Differences in the relative abundance of tRNAs have been linked to modulation of recoding events such as frameshifting (Korniy et al., 2019) and TR (Beznoskova et al., 2016; Blanchet et al., 2014; Roy et al., 2015). Specifically, Trp-, Cys- and Arg-specific tRNAs can be inserted at UGA to stimulate TR (Blanchet et al., 2014; Feng et al., 1990; Urban and Beier, 1995; Zerfass and Beier, 1992). We performed relative quantification of four nc-tRNA isoacceptors that can potentially be incorporated at UGA: Arg-tRNA with the anticodon 5’-UCG-3’ (ArgUCG) (x4) and 5’-UCU-3’ (ArgUCU) (x3), Cys-tRNA with the anticodon 5’-GCA-3’ (CysGCA) (x4) and Trp-tRNA with the anticodon 5’-CCA-3’ (TrpCCA) (x2). In *Drosophila*, each tRNA isoacceptor has several isodecoders, i.e. tRNAs that have different sequences but decode the same codon with varied gene copy numbers; the number of isodecoders for each tRNA isoacceptor is indicated in brackets. The isodecoders for ArgUCU and TrpCCA vary considerably in sequence conservation, which necessitates the use of separate primer pairs for quantification. The isodecoder identity is indicated by a number following the isoacceptor name (eg: TrpCCA1, ArgUCU1,2). Upon quantification, we found that the levels of all three ArgUCU isodecoders were two times more abundant in heads compared to ovaries, while CysGCA and TrpCCA were modestly overexpressed in heads. ArgUCG tRNAs did not show any difference in relative abundance. As a control, we used the non-cognate PheGGA tRNA, which is present in same amounts in heads and ovaries (Figure 6B).

Together, these data indicate the existence of significant differences in the tissue-specific repertoire of eRF1 and eRF3 isoforms, accessory protein factors involved in termination fidelity, and specific nc-tRNAs that can read the UGA stop codon. A combination of these differences can pose a multivariate effect on the selective regulation of termination fidelity in a tissue-specific manner.

## DISCUSSION

TR generates proteins with extended C-termini that can change protein functions and help in adaptation and evolution. A growing body of evidence from bioinformatics and ribosome profiling data indicates that TR is highly pervasive in eukaryotes, but the functions of the extended protein isoforms in an organism remain largely unexplored (Dunn et al., 2013; Jungreis et al., 2016; Jungreis et al., 2011; Williams et al., 2004). Of the 11 candidate genes tested here, several do not undergo TR in S2 cells, which may indicate that some of the predicted candidates are either false positives or do not undergo TR in this particular cell type. High levels of TR in genes ending with UGA-C, such as *svp*, *aPKC, dlg1*, and *tj*, are consistent with the notion that UGA-C is the leakiest stop codon context (Bonetti et al., 1995; Cassan and Rousset, 2001; Cridge et al., 2018; Loughran et al., 2014; McCaughan et al., 1995). The UGA-C sequence can act as the major determinant of TR, as observed with *dlg1* and *tj*. However, in other cases, such as *aPKC*, a regulatory sequence outside of the stop codon context must drive TR. The TR stimulatory elements outside stop codon context can be difficult to predict bioinformatically, but can be of potential use as a synthetic biology tool to induce programmed TR in specific tissues.

Using TR in the *tj* gene as a model, we show that Tj-TR is absent in most tissues, including embryonic and adult gonads, but is selectively expressed in the embryonic CNS and adult brain. The different expression levels are controlled at the translation level, which is particularly surprising, given that the UGA-C termination context appears to be necessary and sufficient for TR in *tj*. Tissue-specific regulation of TR was first reported for the *Drosophila* gene *kelch* (Robinson and Cooley, 1997). Subsequently, ribosome profiling revealed differential ribosomal footprints in several other genes that show significant TR in samples derived from early embryos and S2 cells (Dunn et al., 2013). More recently, several studies in *Drosophila* have reported highest levels of TR in tissues of the CNS, mainly neurons (Chen et al., 2020; Hudson et al., 2020). These studies are concurrent with findings in mice, where the incidence of cell-type-specific TR was detected by analysis of TR in neurons and astrocytes in the CNS as well as peripheral tissues (Palazzo et al., 2020; Sapkota et al., 2019). These observations, combined with significant overrepresentation of neuronal genes in bioinformatically predicted TR candidates (Jungreis et al., 2016; Jungreis et al., 2011) support the idea that elevated TR in susceptible genes might be an idiosyncratic feature of neuronal tissues, albeit the mechanism of such TR regulation remains completely unexplored.

Towards understanding the physiological basis for tissue-specific TR, we found appreciable differences in the expression levels of key players that are directly or indirectly involved in the process of termination. First, we found that the relative abundance of a specific splice variant of *eRF1 (eRF1H)*, is significantly higher in head tissues, comprising 33% of total *eRF1* transcript pool. These isoforms differ in their C-termini. The C-terminal domain of eRF1 is crucial for its interaction with eRF3 (Cheng et al., 2009; Kononenko et al., 2008; Preis et al., 2014). While the depletion of eRF1 has been previously shown to enhance TR at all three stop codons (Carnes et al., 2003; Csibra et al., 2014; Janzen and Geballe, 2004), there are no studies addressing the role of eRF1 isoforms in modulating termination fidelity. Similarly, we also found that the full-length eRF3 isoform is less abundant in head tissues compared to ovaries, while the N-terminally truncated variant of eRF3 (eRF3C) is enriched in the heads. The depletion of full-length eRF3 isoform (eRF3a) is known to increase TR by affecting eRF1 stability in human cell lines (Chauvin et al., 2005; Janzen and Geballe, 2004). A similar eRF3 orthologue termed eRF3b is overexpressed in the brains of mice (Chauvin et al., 2005; Hoshino et al., 1998). eRF3a and eRF3b mainly differ in their N-terminal domains and overexpression of eRF3b, but not of eRF3a, was shown to reduce TR (Jakobsen et al., 2001). So far, it is not known how the C-terminal isoform of eRF1 or the N-terminally truncated eRF3C isoform affect termination fidelity. Together they might be important contributors that lead to selective regulation of TR in nervous tissues. These isoforms are generated by alternative splicing, a phenomenon which is particularly prominent in the nervous system and allows neuronal cells to expand their transcriptomic diversity (Raj and Blencowe, 2015; Su et al., 2018; Venables et al., 2012). Thus, specific cell-types can link several gene regulatory mechanisms that culminate into a regulated expansion of proteome diversity.

Another attractive explanation for tissue-specific TR is the differences in the availability of TR-prone suppressor tRNAs that can read the leaky stop codons due to wobble base pairing at the first or third codon position. Our data indeed support this idea as we found specific tRNAs that are near cognate to UGA, differentially expressed in head and ovarian tissues. Tissue-specific increase in the expression levels of such tRNAs would make them more efficient in competing with release factors for the stop codon, thereby resulting in higher TR. Apart from the core players that directly interact with the stop codon during termination, we also found tissue-specific differences in the levels of *Drosophila* orthologues of PABP, Rli1/ABCE1, Gle1, Dbp5/DDX19 and Pub1. These factors do not play a direct role in stop codon recognition during termination, but are known to have accessory roles that can potentially modulate termination fidelity by interacting with the translational machinery. Taking into account all these findings, it is very likely that the selective elevation of TR in neuronal tissues is caused by a combination of multiple effects. The cellular repertoire that supports TR in susceptible genes is unlikely to disrupt homeostatic cellular proteome, as UAA is the preferred stop codon in highly expressed and housekeeping genes (Trotta, 2016). Additionally, the presence of secondary in-frame stop codons within a few positions downstream of the primary stop codon, or appendage of a peptide sequence that destabilizes the TR extended protein mitigates any dominant negative effects that can be attributed to erroneously extended TR genes (Arribere et al., 2016; Fleming and Cavalcanti, 2019). Thus, selective upregulation of TR in select candidate genes with leaky termination contexts serves to enrich the diversity of the neuronal proteome.

The tight control of TR is biologically important, as constitutive induction of TR in *tj* results in several defective phenotypes in adult ovaries, such as abnormally assembled niches, ambiguous specification of niche cells, ectopic expression of En, deformed germaria, and defective encapsulation of germline cysts by follicular epithelia. Several of these phenotypes are reminiscent of previously characterized hypomorphic *tj* variants (Li et al., 2019; Panchal et al., 2017). Since the expression level of Tj in *tj^TR/TR^* ovaries is comparable to that in *tj^nat/nat^* ovaries, the observed phenotypes can be attributed to altered transcriptional activity caused by the Tj-TR isoform in soma-specific gene regulatory functions during gonad development.

In adult heads, the Tj-TR isoform comprises 20% of the total Tj protein expressed (Figure 2D). Constitutive induction or abolition of TR in *tj* gives rise to several neural phenotypes in the OL and the CB of *tj^TR/TR^* and *tj^nTR/nTR^* mutants. The phenotypes manifest in the formation of sporadic or large multiple lesions in both mutants with comparable frequencies. These results demonstrate that the maintenance of a distinct ratio between the Tj and Tj-TR isoforms is crucial for the preservation of neuronal structure in the brain (Figure 7). The results of the RT-qPCR and RNA-seq experiments begin to uncover a network of potentially complex interactions modulated by Tj-TR. Constitutive expression of Tj-TR leads to upregulation of a number of genes such as *irk3, sand, ninaD*, and *Obp99a*, suggesting that the Tj-TR isoform can act as activator of these targets. The genes that are deregulated upon constitutive induction of TR in *tj* are mostly involved in CNS functions related to perception of external stimuli such as photo stimuli and odorants and regulation of bodily responses towards such stimuli.

**Figure 7.**
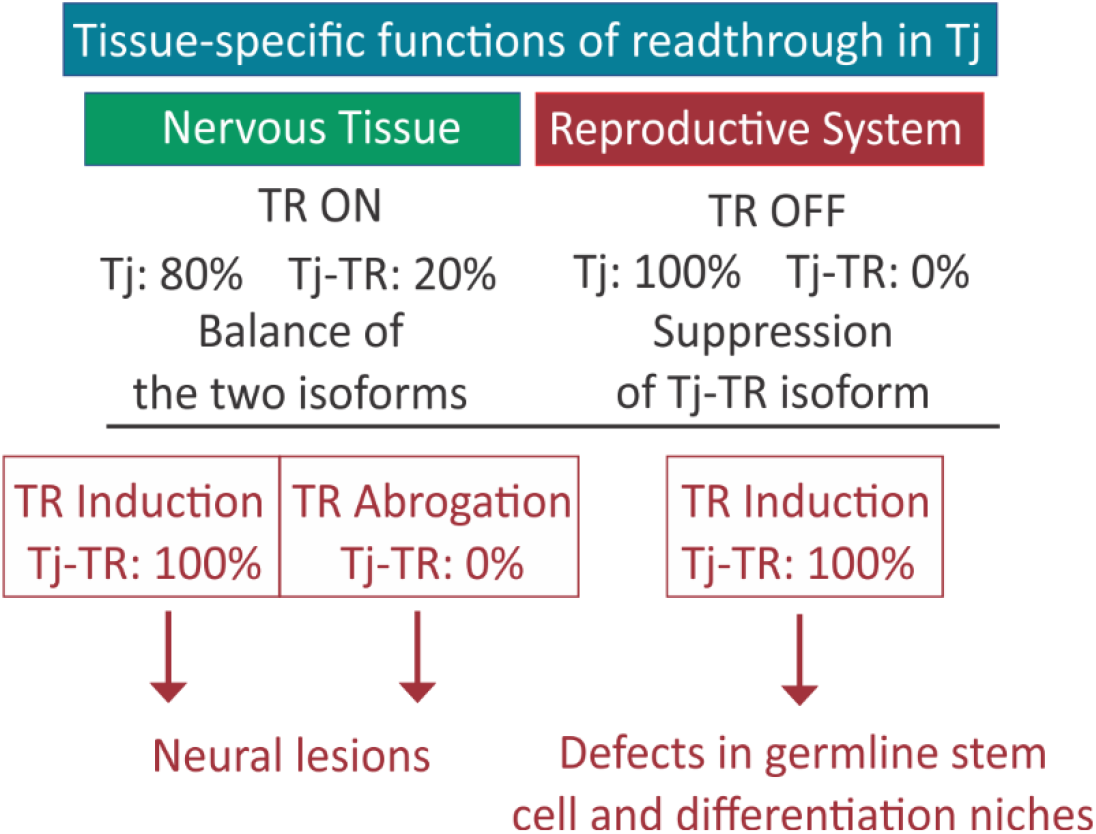
Scheme depicting the importance of proper regulation of Tj-TR. In nervous tissue, TR of *tj* transcript results in a mixture of Tj and Tj-TR isoforms. Abrogation of either isoform results in neural lesions, while a properly regulated balance of isoforms is key to brain health. In the reproductive system, preventing Tj-TR is crucial for reproductive capacity.

Because the function of Tj in the nervous system is not well understood, the exact mechanism of regulation of Tj function by TR is not clear at present. TR in Tj does not change the protein localization in the cell, unlike with those proteins where the peptide added by the TR contains a nuclear localization signal or a transmembrane domain (Bersch et al., 2018; Dunn et al., 2013). Given the proximity of the TR extension to the DNA binding leucine zipper motif in the C-terminal region of Tj, it is possible that the addition of the TR motif impedes existing or confers novel DNA binding properties, or that the TR motif alters the protein-binding properties of Tj. For example, the TR motif in AQP4 appends a phosphorylation site and is important for interaction with other proteins (De Bellis et al., 2017). Furthermore, bZIP transcription factors, of which Tj is an example, are known to form homo/heterodimers using their dimerization domains (Kataoka et al., 1994; Kurokawa et al., 2009). The TR extension might affect the dimerization of Tj, which would lead to attenuation of gene regulatory functions of the native Tj. The structurally disordered C-termini generated by TR (Kleppe and Bornberg-Bauer, 2018; Pancsa et al., 2016) may ensure the accessibility for new interactors or affect liquid-liquid phase transitions involving transcription factors in the nucleus (A and Weber, 2019; Owen and Shewmaker, 2019), thereby affecting the native protein function. Given the high number of TR genes involved in regulatory functions in *Drosophila*, an interaction-prone C-terminal segment would allow fine-tuning of the protein function in a regulated manner.

The Tj-TR isoform is upregulated upon exposure to selective stress conditions such as heat and oxidative stress. Effector genes that mediate stress responses are controlled by pleiotropic regulatory cascades; Tj-TR might be part of an integral mechanism that allows flies to adapt to stressful environments. Because Maf transcription factors are involved in the regulation of a large number of genes, and Tj is the only large Maf transcription factor in *Drosophila*, TR might allow rapid fine-tuning to modulate CNS responses to the external environment. Alternatively, enhanced TR might also be a by-product of altered translational fidelity due to changes in the components of translational machinery upon exposure to stress, such as starvation or oxidative damage. For example, stress induction by serum starvation enhances drug-induced TR in colorectal cancer cell lines as well as reporter cell lines (Wittehstein et al., 2019). In any case, TR in *tj* links the transcriptional and translational control to fine-tune the gene expression in selected cells.

Neuron-specific genes are highly enriched in the phylogenetically predicted list of TR candidates in *Drosophila* (Jungreis et al., 2011). The nervous system plays an important role in modulating organismal response to various stress conditions. Stress can arise externally due to environmental fluctuations in light, temperature, oxygen levels, and nutrient abundance or internally due to molecular damage during development. A rapid response to such cues not only restores homeostatic equilibrium but also ensures survival. Such responses include regulating the expression of a battery of transcription factors and other molecules (miRNAs, enhancers, etc). In such a scenario, tweaking the neuronal proteomic diversity by post-transcriptional or translational regulatory mechanisms would confer selective advantage by generating a rapid response. For example, adenosine-to-inosine (A-to-I) editing, which is most active in the nervous system, is an important contributor to neuronal dynamics (Behm and Ohman 2016). TR might represent yet another important mechanism to fine tune neuronal gene expression and create protein isoforms with altered properties. Exploring the repertoire of TR-generated proteins might significantly augment our understanding of the neuronal proteomic diversity and its implications on physiological processes.

## STAR*METHODS

### Key resource table

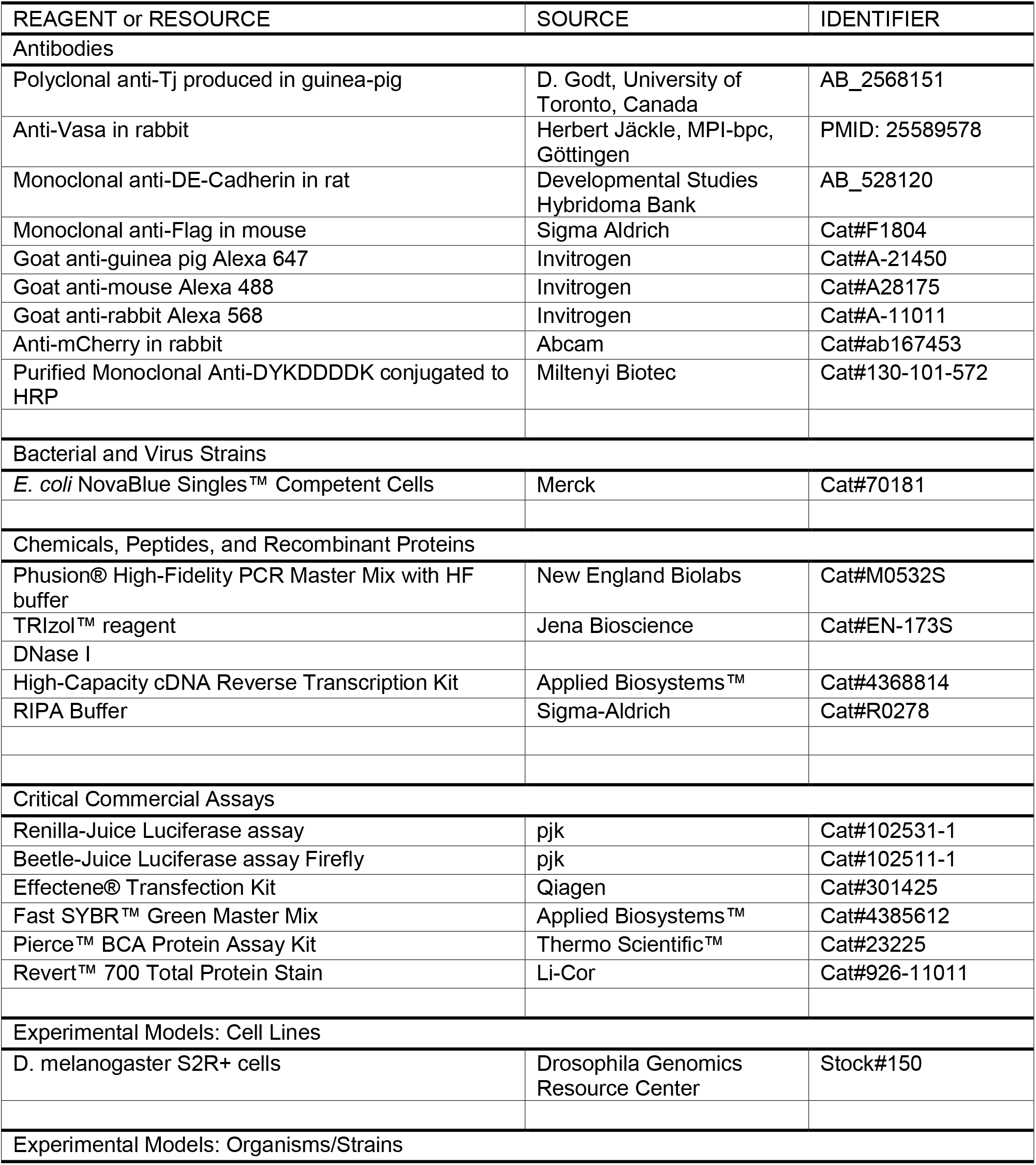
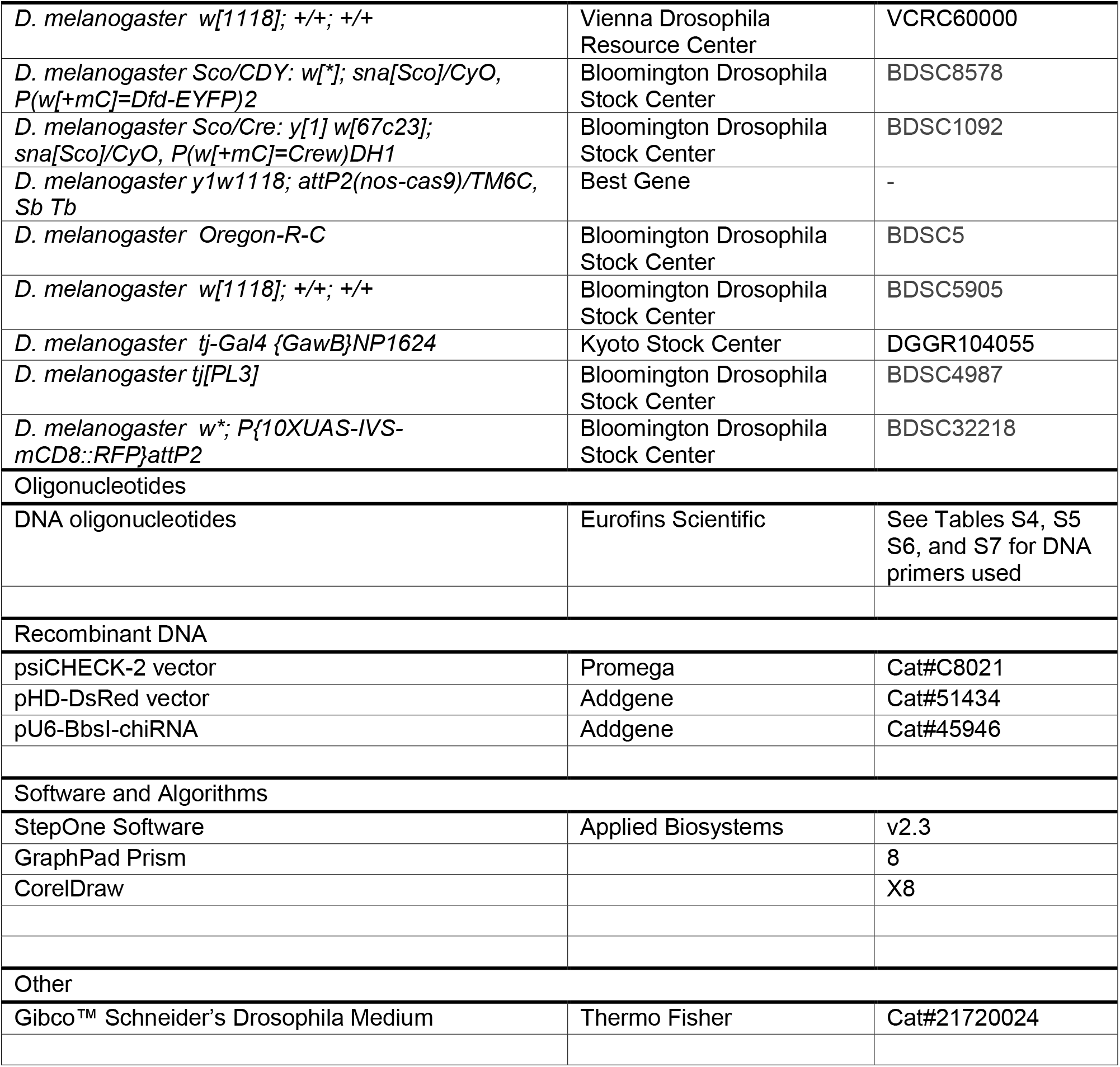

## Experimental model and subject details

### *E. coli* cells

*E. coli* NovaBlue Singles^™^ Competent cells-Novagen *endA1 hsdR17* (r_K12^-^_ m_K12^+^_) *supE44 thi-1 recA1 gyrA96 relA1 lac* F’[*proA^+^B^+^ lacI^q^Z*Δ*M15*::Tn*10*](Tet^R^) were used for all cloning and plasmid construction purposes. 50 μl of chemically competent cells were used for transformation. Thawed cells were transformed with 1-5 μl of PCR, ligation, or Gibson assembly products on ice for 20 min. The cells were heat shocked by incubating at 42 °C for 45 s. The transformed cells were kept on ice for 2 min, supplemented with 450 μl LB media and incubated at 37 °C for 1 hour. 100 μl of the culture was plated on LB agar containing the appropriate selection antibiotic. The colonies were grown 37 °C in LB media with appropriate antibiotics and screened via plasmid DNA sequencing (Microsynth AG).

### Schneider 2 cells

*Drosophila* S2R+ cells were cultured in 25 cm^2^ flasks at 25°C in a CO2 incubator in Schneider’s *Drosophila* medium (Gibco^®^), supplemented with 10% heat-inactivated fetal bovine serum, 100 units/ml penicillin, and 100 μg/ml streptomycin. Cells were passaged every 3-4 days by splitting them in 1:6 ratio. Prior to transfection, S2R+ cells were split 1:6 and 150 μl of the cell suspension was seeded into flat-bottomed 96-well plates and incubated overnight. Transfection reactions were prepared using Effectene^®^ Transfection Kit. 100 ng of dual reporter construct was used to transfect each well, when the cell confluency reached approximately 70%. The cells were incubated for 72 hours to allow luciferase expression. Transfections were performed in triplicates.

### *Drosophila* lines

Fly stocks were maintained on standard food with yeast, cornmeal, and agar in a controlled environment with constant temperature of 25 °C, constant humidity, and 12 hr-12 hr light-dark cycle, unless otherwise stated. For control animals, we collected the progeny of *w^1118^* males crossed to *OregonR* virgin females.

## Method details

### Plasmid construction

Vector templates and/or inserts for molecular cloning were amplified by PCR using Phusion High-Fidelity PCR master mix. Desired PCR products were purified using Nucleospin^®^ Gel and PCR clean up kit. Point mutations were introduced into plasmid vectors using the Agilent QuikChange II site-directed mutagenesis protocol. Insertional and deletion mutagenesis were performed using a blunt-end ligation method. Phosphorylation and ligation reactions were performed using T4 Polynucleotide kinase and T4 ligase, and the ligated products were transformed into competent cells. Molecular cloning of TR test cassettes into psiCHECK^™^-2 vector, HA and TfR into pHD-DsRed was achieved by isothermal assembly (Gibson assembly) (Gibson et al., 2009). Insert sequences for Gibson assembly were amplified from gDNA or cDNA using primers containing 18 bp overhangs that overlap with the blunt ends of PCR-amplified, linearized vectors. Gibson assembly was performed by incubating the purified inserts and 100 ng linearized vectors in a molar ratio of 3:1 with ‘in-house’ prepared Gibson assembly mix for 1 h at 50 °C in a total volume of 15 μl. 1 μl of end product was transformed into competent cells.

### Dual Luciferase reporter assay

The commercial psiCHECK^™^-2 (Promega) vector was modified by deleting the Rluc poly(A) signal and the Fluc HSV TK promoter using blunt-end ligation, such that both the reporters are transcribed as a single transcriptional unit from a monocistronic mRNA controlled by an SV40 promoter. A self-cleaving P2A sequence (66 bp) was inserted between the two reporters using two-step insertional blunt-end ligation mutagenesis. The start codon of Fluc was deleted using blunt-end ligation mutagenesis. The primers used for vector modification are listed in Table S5. TR test cassettes (105 bp) from the candidate genes containing the leaky stop codons, flanked by 51 bp on either side, were amplified from *w^1118^* cDNA using primers with 18 bp overhangs and inserted into modified linearized vector using Gibson assembly. UGA to UUC and UAAA point mutations were introduced into each construct using blunt-end ligation mutagenesis. The primers for cloning of TR motifs and point mutations are listed in Table S6.

The activities of Renilla and Firefly luciferases were measured 72 hours after transfection using Beetle juice and Renilla Glow Juice. Measurements were performed in a luminometer with a delay time of 2 seconds and an integration time of 5 seconds. Background luminescence (obtained from cell lysates prepared from S2 cells transfected with empty transfection mixes) was subtracted from the raw readouts of the luminescence signals. The ratio of Fluc:Rluc signal was calculated for each construct containing the native and the UAA-A stop codon context in their TR motif, as well as for the corresponding constructs where the stop codon is mutated to UUC. To calculate TR efficiency of test constructs with native and UAA-A stop codon contexts, their respective Fluc:Rluc values were divided with Fluc:Rluc values of constructs containing a UUC codon, which serve as a positive controls. Non-paired two-tailed Student’s t-test was used to analyze the results.

### CRISPR/Cas9 design

CRISPR/Cas9-based genome editing was employed to create three different genetic mutants of *D. melanogaster* that harbor mutations in and around the stop codon of the *traffic jam* (*tj*) gene (sequence location 2L:19,64,267 to 19,467,758). The CRISPR target finder tool (http://tools.flycrispr.molbio.wisc.edu/targetFinder/) was used to find optimal PAM sites on the *tj* gene that flank the TR region between the first and the second stop codon of the *tj* ORF. The proximal PAM site was 5’ AGAGCTTT|GGCTATCGCCGC **CGG** 3’ and the distal PAM site was 5’ ACACAATG|TATAAGGTAAAT **TGG** 3’, where the NGG motifs are highlighted in bold. The 20 bp proximal and distal PAM regions were introduced upstream of gRNA scaffold into two separate pU6-BbsI-chiRNA vectors (Addgene) via blunt-end ligation-mediated insertional mutagenesis using primer pairs PK241_F/PK243_R and PK242_F/PK243_R respectively (Table S7).

pHD-DsRed vector (Addgene) carrying the homology arm 1 (HA1), the Template for Recombination (TfR), and the homology arm 2 (HA2) were generated in subsequent steps using Gibson assembly. HA1 (1100 bp) + TfR (250 bp) was amplified from gDNA obtained from *w^1118^* as a single fragment and inserted upstream of loxP-DsRed-SV40poly(A)-loxP sequence. HA2 (1144 bp) was amplified and inserted immediately downstream of this sequence. QuikChange mutagenesis protocol was used to introduce synonymous mutations into the proximal PAM sequence that borders HA1 and TfR, in order to prevent Cas9 from cleaving the vector once injected into the embryos. UGA to UUC mutation was then introduced in the TfR at the *tj* stop codon by QuikChange mutagenesis and 3xUAA was inserted downstream of the *tj* stop codon by blunt-end ligation method. 3xFlag was inserted upstream of the second stop codon by Gibson assembly. These cloning steps were performed in a pHD-DsRed vector in which the loxP1 site had been deleted in order to avoid complications associated with redundant primer binding sites. Finally, the loxP1 site was reinserted. Additionally, the dispensable phage pC31 attP site was removed from the pHD-DsRed vector during PCR amplification. The primers used for Gibson assembly, point mutations, and blunt-end ligation cloning are listed in Table S7. Due to the introduction of an independent SV40 transcription termination signal in the TfR, the biogenesis of *tj*-derived piRNAs in the CRISPR-derived recombinants is inhibited. To overcome this limitation, the loxP-flanked *DsRed*-SV40 poly(A) marker cassette was removed by Cre-Lox recombination, which restored the native *tj* 3’ UTR in *tj*-TR mutants (*tj^mut^*). The introduction of the desired mutations was verified via sequencing of a genomic DNA-derived amplicon. *DsRed* deletion was confirmed via screening of eyes for negative fluorescence as well as by sequencing. RH genome engineering services offered by Best Gene Inc. Chino Hills, CA, USA were used for CRISPR injections. The fly strain used for injection had the following genotype: *y^1^w^1118^*; *attP2(nos-cas9)/TM6C, Sb Tb*.

DsRed-positive CRISPR mutants were crossed with *Sco/CDY* balancer lines to obtain CDY-balanced mutant lines for second chromosome. *tj^mut^*(+DsRed)/*CDY* lines were crossed with *Sco/Cre* lines in order to achieve Cre recombinase-mediated removal of the DsRed marker. The progenies, *tj^mut^*(±DsRed)/*Cre*, were back crossed with *Sco/CDY* balancer lines to obtain DsRed-deleted, *tj^mut^*/*CDY* flies that served as stocks. DsRed deletion was confirmed by screening individual balanced flies for the absence of DsRed. The *tj^mut^/tj^mut^* obtained by back crossing of *tj^mut^/CDY* flies were used for experimental purposes. gDNA was extracted from the homozygous mutant flies. Using it as template, the genomic region flanking PAM sites was amplified using primers PK277_F and PK278_R and sequenced using primers PK277_F and PK279_F to confirm the introduced mutations (Table S7). *w^1118^* flies were used as wild-type controls as they have the closest genetic background to the mutants.

### Immunohistochemistry

Immunofluorescent studies were performed using standard procedures (König and Shcherbata, 2013). Primary antibodies used were: guinea pig anti-Tj (1:10,000; a kind gift from Dorothea Godt, University of Toronto, Canada), rabbit anti-Vasa (1:5,000; a gift from Herbert Jäckle, MPI-bpc, Göttingen), mouse anti-Engrailed (1:20; Developmental Studies Hybridoma Bank, DSHB), mouse anti-Fas3 (1:20; DSHB), rat anti-DE-Cadherin (1:20; DSHB), rabbit anti-mCherry (1:200; Abcam), and mouse anti-Flag (1:500; Sigma Aldrich). The following secondary antibodies were used: goat anti-guinea pig Alexa 647 (1:500; Life Technologies, A-21450), goat anti-mouse Alexa 488 (1:500; Molecular Probes) and goat anti-rabbit Alexa 568 (1:500; Molecular Probes). DAPI (Sigma) was used to stain the nuclei. Samples were imaged using Zeiss LSM 700. ImageJ and CoreDrawX8 were used to make figures.

### Histology of *Drosophila* brains

For analysis of adult brain morphology, 7 μm paraffin-embedded sections were cut from fly heads. To prepare *Drosophila* brain sections, the fly heads were immobilized in collars in the required orientation and fixed in Carnoy fixative solution (6:3:1 = Ethanol:Chloroform:Acetic acid) at 4°C overnight. Tissue dehydration and embedding in paraffin was performed as described previously (Kucherenko et al., 2010). Histological sections were prepared using a Hyrax M25 (Zeiss) microtome and stained with hematoxylin and eosin as described previously (Shcherbata et al., 2007). All chemicals for these procedures were obtained from Sigma Aldrich.

### Gene expression analysis

Standard miniprep protocol for *D. melanogaster* genomic DNA extraction was followed as described in Huang et al. (2009). Total RNA was extracted from heads and ovaries of 3-4 days old adult flies of each genotype using TRIzol reagent following manufacturer’s protocol. Extracted RNA was quantified and treated with DNaseI (2 units per μg of RNA). Total cDNA was prepared using random primers with High Capacity Reverse Transcriptase (ThermoFisher) following manufacturer’s instructions. 20 μl RT reactions were set up for 1 μg RNA template. RT-qPCR was performed using Quantitect SYBR Green PCR kit. Each reaction was performed in 15 μl volume using 20 ng cDNA template and 200 nM primers. All reactions were performed in triplicates. Control reactions were set up for each target gene using non-RT templates. The primers used for each transcript quantification were obtained from DRSC primer bank and are listed in Table S8. The qPCR reaction conditions used were according to manufacturer’s instructions. *αTub84B* was used as endogenous control and *tj^nat/nat^* flies were used as control samples. The analysis of the acquired threshold cycle (CT) values was performed using StepOne Software. CT value denotes the fractional cycle number at which the fluorescence signal for each test sample passes a defined threshold. Average C_T_ values from three technical replicates of respective genes were subtracted from that of the *αTub84B* control to obtain ΔC_T_. These ΔC_T_ values for each gene was normalized again by subtracting the ΔC_T_ of the control sample from the ΔC_T_ of the test sample. The ΔΔCT values thus obtained were used to calculate gene expression levels by using the formula RQ = 2^-ΔΔCT^. Non-paired two-tailed Student’s t-test was used for calculating p values. For quantification of individual cellular tRNAs, a modified stem-loop qPCR method was used as published by (Wan Makhtar et al., 2017). Stem-loop reverse transcription primers were used for individual tRNAs (Table S9). Separate stem-loop primer was also designed for 18S rRNA which is used as an internal control. cDNA was prepared using SuperScript III^™^ Reverse Transcriptase following manufacturer’s protocol. The amplification for individual tRNAs using RT-qPCR was performed using tRNA-specific forward primers and a universal reverse primer (Table S9). RNA sequencing services were provided by Transcriptome and Genome Analysis Laboratory (TAL), Göttingen.

### Stress induction and fly survival assays

Stress induction experiments were carried out with 5-6 days-old flies. Approx. 50 flies were used for each condition. Flies were starved by placing them in empty vials with glass fiber disk soaked in water. Acute heat stress was induced by incubating the flies at 36 °C for 3 hours. Oxidative stress was induced by placing flies in vials with glass fiber disk soaked in 5% sucrose and 20mM paraquat solution for 12 hours. For sleep deprivation, vials containing flies were shaken at 300 rpm under constant light source for 24 hours. To study ageing, flies were reared on standard food vials for 3 weeks, with food change every week. The method for lysate preparation from stressed fly heads is described in the following section. The survivability rate of flies upon oxidative stress was recorded by observing fly activity for 3 days upon feeding them 5% sucrose, 20 mM paraquat solution as mentioned above. 20 flies were taken for each genotype and the study was performed in five biological replicates.

### Quantitative Western Blot

Whole tissue lysates were prepared from ovaries and heads using RIPA buffer (Sigma Aldrich), and the protein concentration was measured using Pierce^™^ BCA Protein Assay Kit. Equal amounts of total protein extract were loaded as samples for western blot. Instead of using housekeeping genes as loading controls, we utilized the total protein normalization (TNP) method (Licor 700 Total Protein Stain). The TNP staining gives a linear signal range, which is especially relevant for low-abundance proteins such as Tj. Anti-DYKDDDDK-HRP antibodies (Miltenyi Biotec) were used to immunostain Tj-TR-3xFlag bands. HRP signal was developed using Chemiluminescent substrates (ThermoFisher) and imaged in the Odyssey Fc Imaging System. The anti-Flag chemiluminescent signal from each tissue derived from *tj^nat/nat^* mutants was normalized to the respective TNP signal and compared to the anti-Flag/TNP signal from tissues derived from *tj^TR/TR^* mutants to calculate TR efficiency of Tj.

### Quantification and Statistical Analysis

For luciferase assays, error bars show the standard deviation obtained from at least three technical replicates (n=3). For constructs with stop codon context of UGAC, three independent biological replicates were performed. Statistical significance was determined by two-tailed unpaired Student’s t-test wherein p-values of < 0.05 were considered significant in all cases. RNAfold web server was used for prediction of secondary structures on mRNA sequences. All western blots were performed using head samples from three biological replicates for each mutant fly line. For quantification of niche and germarium phenotypes, n>120; for quantification of neural lesions in central brain, n~25 and in optic lobe, n~50 were used. Two-way tables and Chi-square testes were used to determine significant differences in the phenotypes between the mutants. Statistical analyses were performed with Graphpad prism 8. RT-qPCR studies were performed from cDNA obtained from three independent biological samples. Three technical replicates were used for each biological sample. StepOne Software v2.3 was used to perform comparative C_T_ (ΔΔC_T_) quantification. The data were normalized against average ΔC_T_ of the housekeeping gene *αTub84B*. Data values from tjnat/nat sample were used as endogenous control. p-values are calculated using two-tailed unpaired Student’s t-test with Welch’s correction of standard deviation from ΔΔC_T_ values of *tj^nat/nat^* and *tj^TR/TR^* samples.

For quantification of neural lesions, central brains and optic lobes were analyzed independently. For immunostained brains, z-stack images were acquired through the brains using a Zeiss LSM700 confocal microscope. Examination of all slices was taken into account to identify lesions, since they are found to occur throughout both CBs and OLs of analysed brains. For H&E-stained and sectioned brains, multiple sections for each brain were viewed and assessed for presence of lesions. In both cases, a distinction was made between the presence of multiple large lesions and sporadic lesions or a sponge-like appearance of the brain tissue.

For quantification of niche, germarium, and ovariole phenotypes, we used immunofluorescent staining for common markers to discern the presence of mutant phenotypes. Ovaries from each genotype were stained, and then ovarioles were separated from one another on slides prior to imaging. Each ovariole, corresponding germarium, and GSC niche was examined, and each was counted independently as either normal or exhibiting mutant phenotypes. En staining was used to evaluate GSC niches, as cap cells are countable with this marker. Vasa staining was used to assess the health and differentiation of the germline. Fas3 staining was used to visualize the follicular epithelium and to monitor the process of germline encapsulation in the posterior end of germaria. Finally, DAPI combined with all other stains was used to evaluate the overall shape and size of germaria; in addition, condensed and fragmented DAPI staining was used as an indicator of cell death in egg chambers. Two-way tables and Chi-square tests were used to determine significant differences in the phenotypes between the mutants.

## ACKNOWLEDGMENTS

We thank Jasmin Rehman and Ibrahim Ömer Cicek for their invaluable scientific insights. We thank Anna Pfeifer, Olaf Geintzer, Sandra Kappler, Christina Kothe, Theresia Niese, Tanja Wiles, Vanessa Herold, Franziska Hummel, Tessa Hübner, and Michael Zimmermann for expert technical assistance. The work was supported by a grant of the Deutsche Forschungsgemeinschaft (SFB860 to M.V.R.)

## AUTHOR CONTRIBUTIONS

P.K. prepared materials. P.K., T.D.C., A.S.Y. and H.R.S. performed experiments and analyzed the data; all authors conceived the research, discussed the results and wrote the manuscript.

## DECLARATION OF INTERESTS

The authors declare no competing interests

## Supplemental Information

**Figure S1.**
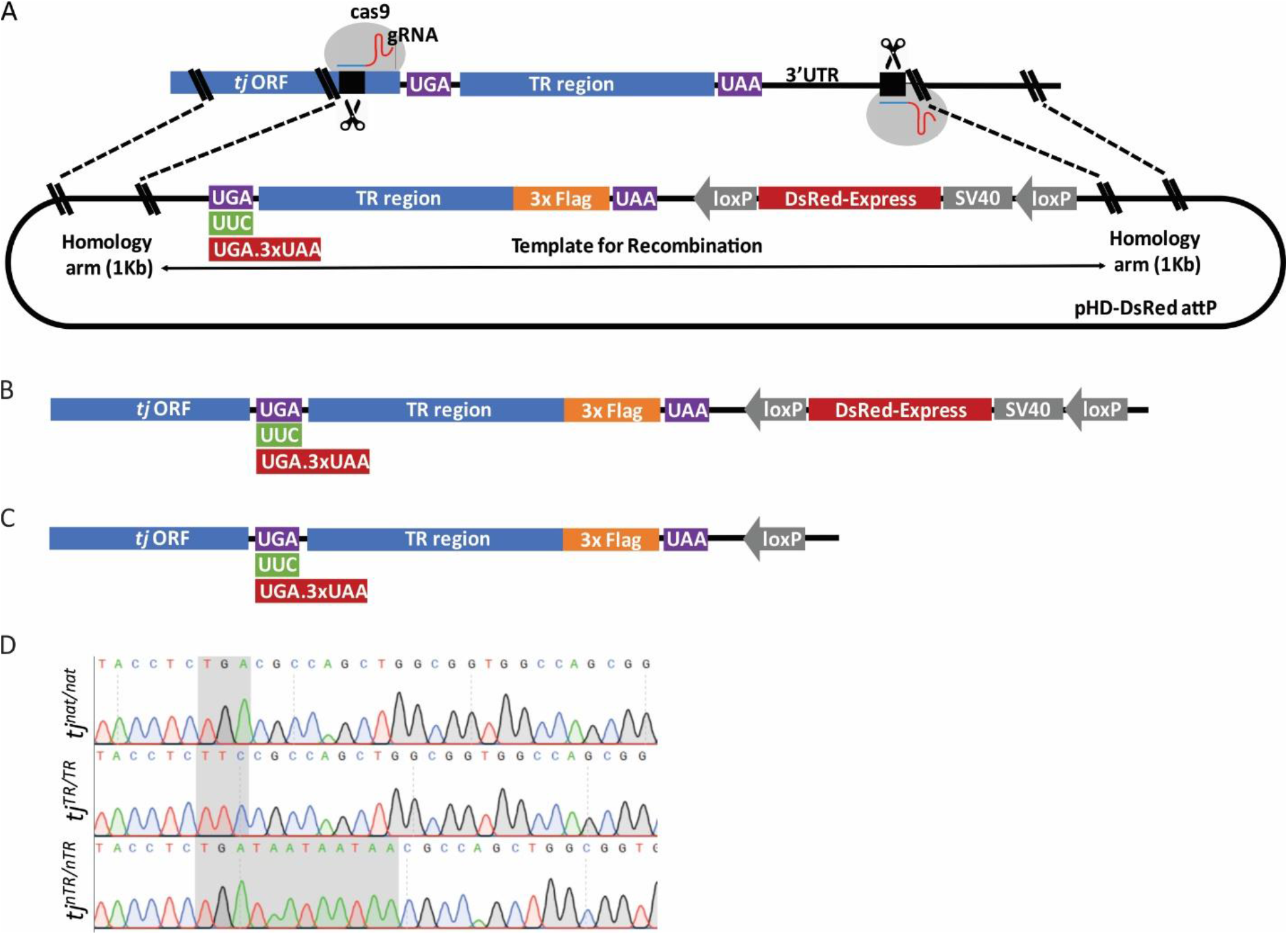
Construct design for CRISPR/Cas9-mediated genome editing to create *tj*-TR mutants, Related to Figure 2. (A) Gene locus surrounding the TR region of *tj* with proximal and distal PAM (protospacer adjacent motif) sites for guide RNA (gRNA)-directed Cas9 cleavage depicted above the modified pHD-DsRed attP vector containing the Template for Recombination (TfR) flanked by 1-Kb homology arms. Dotted lines represent the region of homology between the gene locus and modified vector. The TfR contains the modifications that introduce the desired mutations at the primary *tj* stop codon, 3xFlag upstream of the second stop codon and a loxP flanked *DsRed* marker. (B) Sequence depicting the modifications introduced in the *tj* locus in *tj*-TR mutants post-CRISPR/Cas9 editing. (C) Restoration of the native 3’ UTR in *tj*-TR mutants via Cre recombinase-mediated removal of the loxP-flanked DsRed marker. (D) Sequence verification of *tj^nat^, tj^TR^ and tj^nTR^* mutations in homozygous mutant flies.

**Figure S2.**
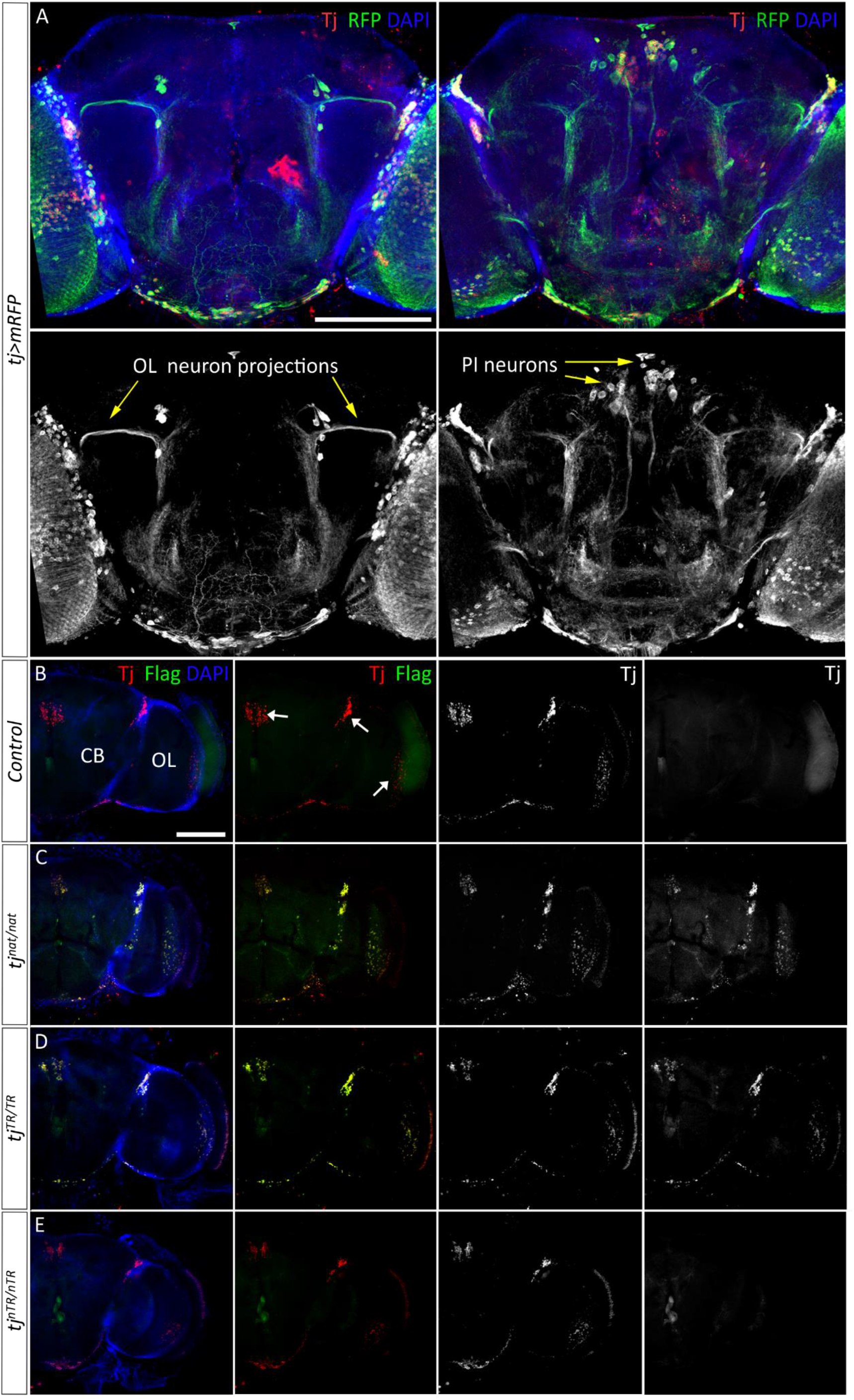
Visualization of Tj and Tj-TR isoforms in adult brains from *tj*-TR mutants, Related to Figures 2 and 3. (A) Membrane-bound RFP was expressed under the control of *tj-Gal4*, an enhancer trap that closely mimics the endogenous expression pattern of *tj*. RFP marks cell bodies as well as neuronal projections of *tj*-expressing cells. Left panels depict projection patterns in an anterior slice from a z-stack of a whole brain, and the right panels depict a more posterior slice (RFP alone is shown in greyscale panels). Many cells of the retina and optic lobe (OL) express RFP, as do a smaller number of central brain (CB) neurons. In particular, OL neuronal projections extend into the CB (left panes), and a subset of *pars intercerebralis* (PI) neurons are visible near the brain midline (right panels). (B)-(E) Each row depicts one brain hemisphere with a medial CB to the left and the lateral OL to the right. (B) Control brain expressing Tj protein in several clusters in both the CB and the OL. (C) *tj^nat/nat^* brains expressing native Tj in a pattern indistinguishable from control as well as the Flag-tagged TR isoform, demonstrating the occurrence of TR in these cells. (D) Overlapping pattern of Tj and Flag staining in *tj^TR/TR^* brains consistent with constitutive induction of TR in Tj. (E) Abrogation of TR in Tj observed in *tj^nTR/nTR^* as evidenced by the lack of Flag stain. Scale bars: 100 μm.

**Figure S3.**
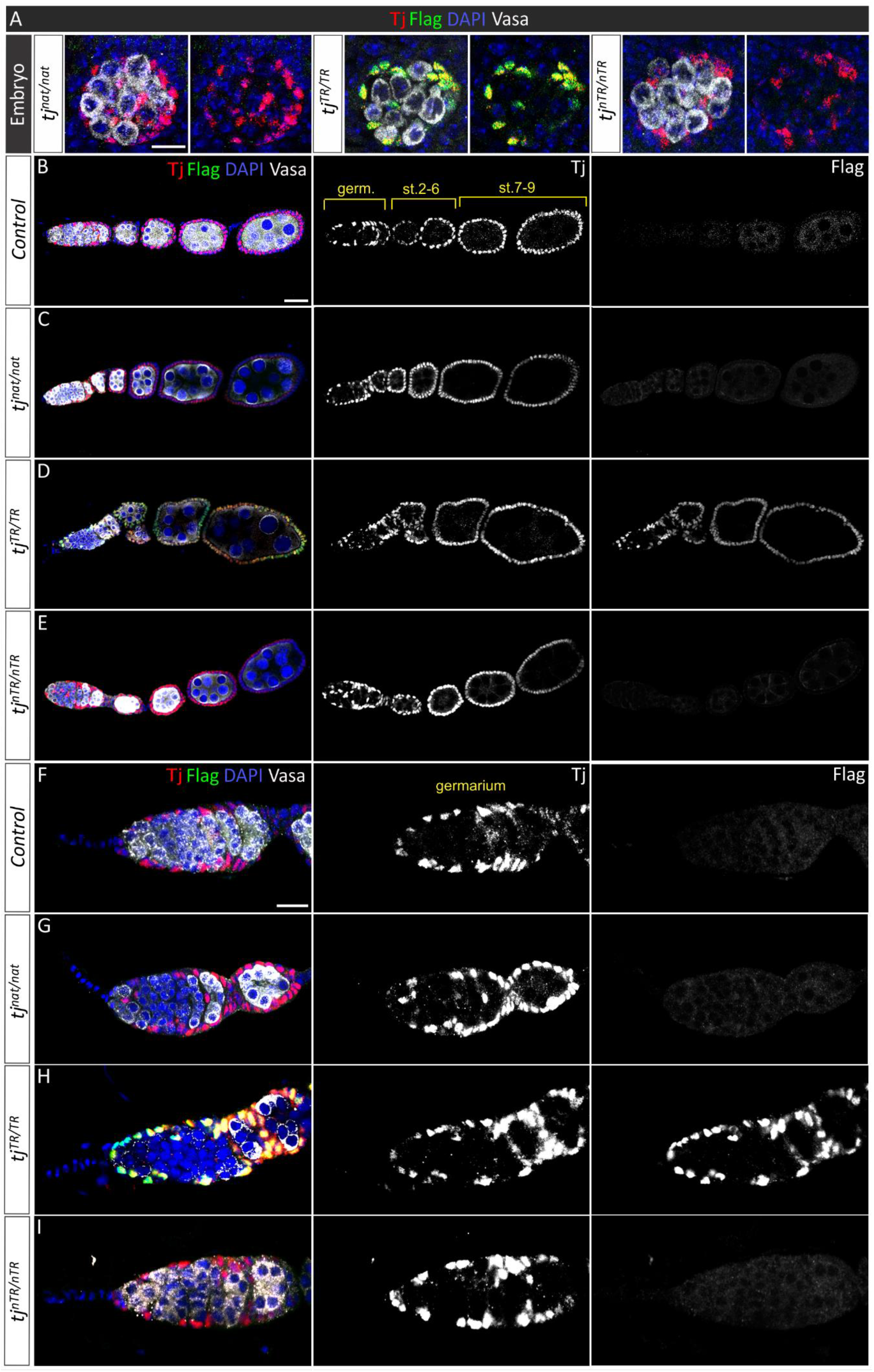
Repression of TR in Tj in adult ovaries, Related to Figures 2 and 4. (A) Embryonic gonads. In all mutants, Tj is expressed in somatic gonadal precursors, not in germline cells (marked with Vasa). Overlap of Tj and Flag staining with nuclear DAPI stain demonstrates that both Tj and Tj-TR are nuclear. In *tj^nat/nat^* and *tj^nTR/nTR^* gonads, lack of Flag stain shows that TR does not occur in these mutants. In contrast, in *tj^TR/TR^* gonads, there is perfect overlap of Tj and Flag. Scale bar: 5 μm. (B-E) Anterior segment of ovarioles depicting most of the endogenous Tj protein expressed in control and in *tj*-TR mutants. In addition to the germarium, Tj expression is robust in follicular epithelia (B). Lack of Flag staining in the *tj^nat/nat^* ovariole (C) demonstrates that Tj-TR does not occur at detectable levels in adult ovaries. *tj^TR/TR^* ovariole (D) shows constitutive expression of Flag-tagged Tj-TR isoform that co-localizes with native Tj. *tj^nTR/nTR^* ovariole (E) does not express the Tj-TR isoform. Scale bar: 20 μm. (F-I) Expression of Tj and Tj-TR isoforms in control and mutant germaria. In controls, Tj is present in follicle cells in the posterior end of each germarium as well as in escort cells and cap cells (F). As observed in (B-D), the *tj^nat/nat^* germarium (G) and the *tj^nTR/nTR^* germaria (I) do not express the Flag-tagged Tj-TR isoform. In contrast, the *tj^TR/TR^* germarium (H) exhibits constitutive co-localization of Tj and Tj-TR isoforms. Scale bar: 10 μm.

**Figure S4.**
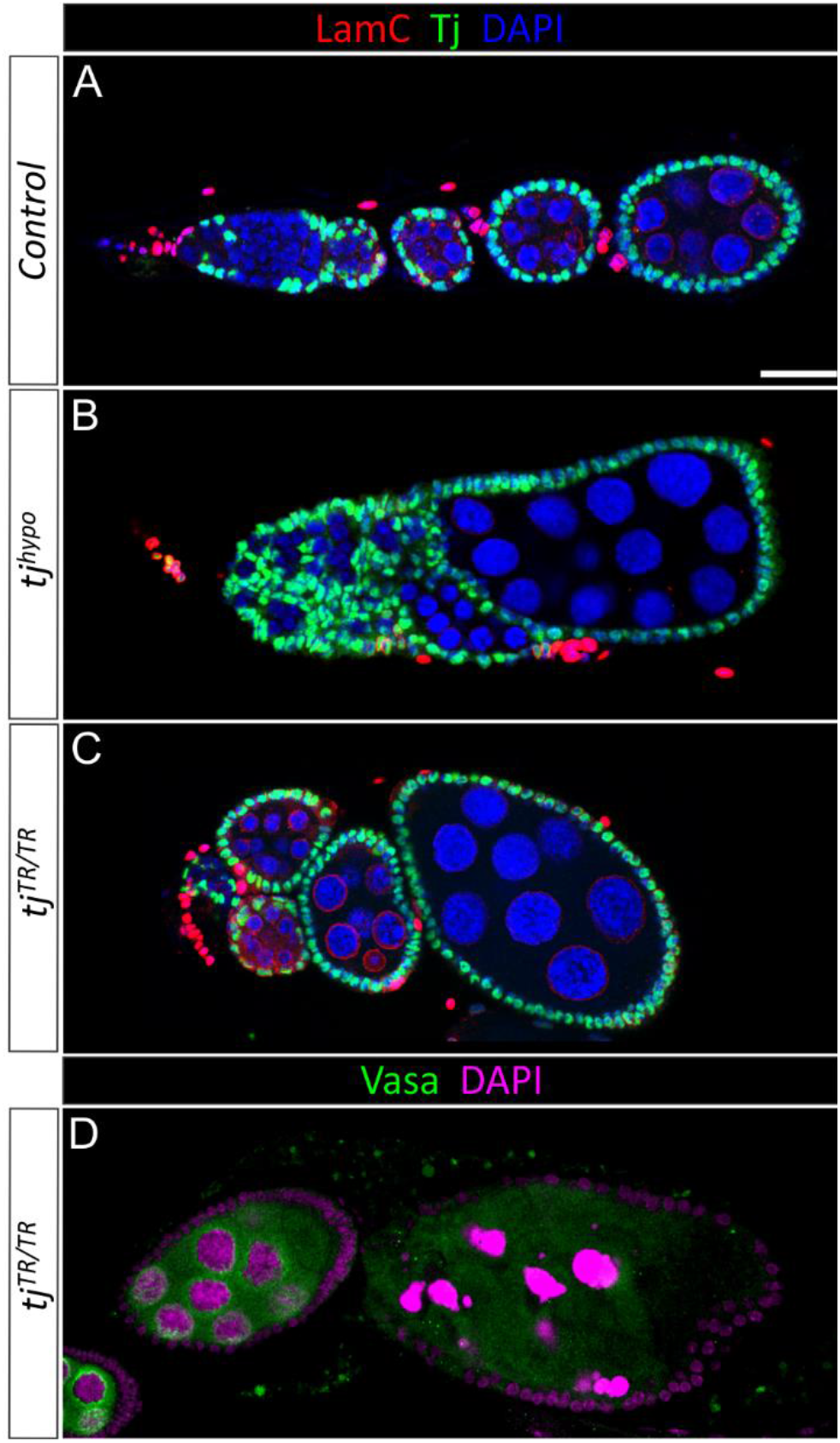
Forced *tj* TR causes various strong ovarian phenotypes similar to those in *tj* hypomorphic ovaries. (A) Control ovariole stained for the nuclear marker DAPI as well as anti-Tj and anti-LamC. Anti-LamC is a nuclear envelope marker that strongly stains TFCs and CpCs as well as stalk cells (the cells that connect adjacent egg chambers) and more weakly marks nurse cells. It is also visible in the nuclei of the muscle cells that surround each ovariole. (B) A *tj* hypomorphic ovariole exhibiting a strong disorganization phenotype. The egg chambers are not properly separated, and the germarium is very small or absent. Note that a LamC-positive TF is still visible at the anterior tip (to the left of the panel). (C) A *tj^TR/TR^* germarium with a very small germarium. Similar to the hypomorph, the TF is still present. (D) A *tj^TR/TR^* germarium exhibiting cell death in the germline of the larger egg chamber, evidenced by the strongly condensed and fragmented DAPI staining. Scale bar: 20 μm.

**Figure S5.**
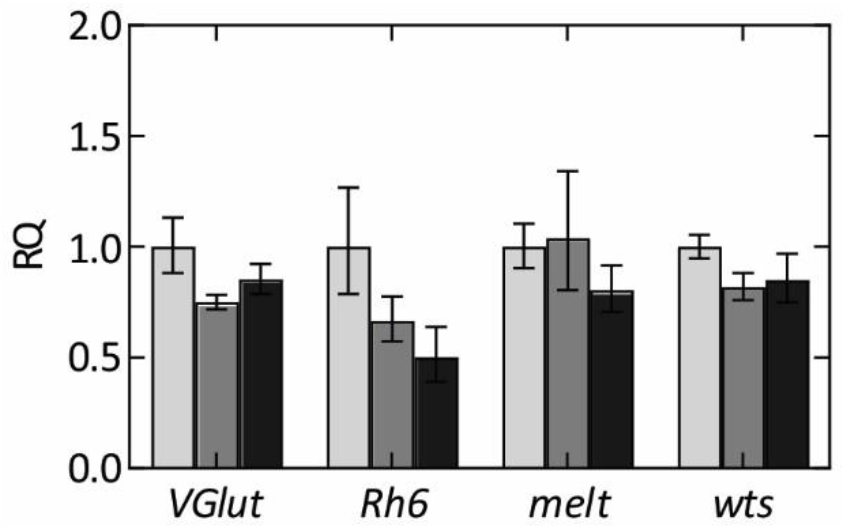
RT-qPCR analysis of genes known to be regulated by Tj in *tj*-TR mutants using cDNA prepared from adult heads. Error bars represent the upper and lower limits of RQ values defined by the standard deviation of ΔΔC_T_. The data were normalized against average ΔC_T_ the housekeeping gene *αTub84B*.

**Figure S6.**
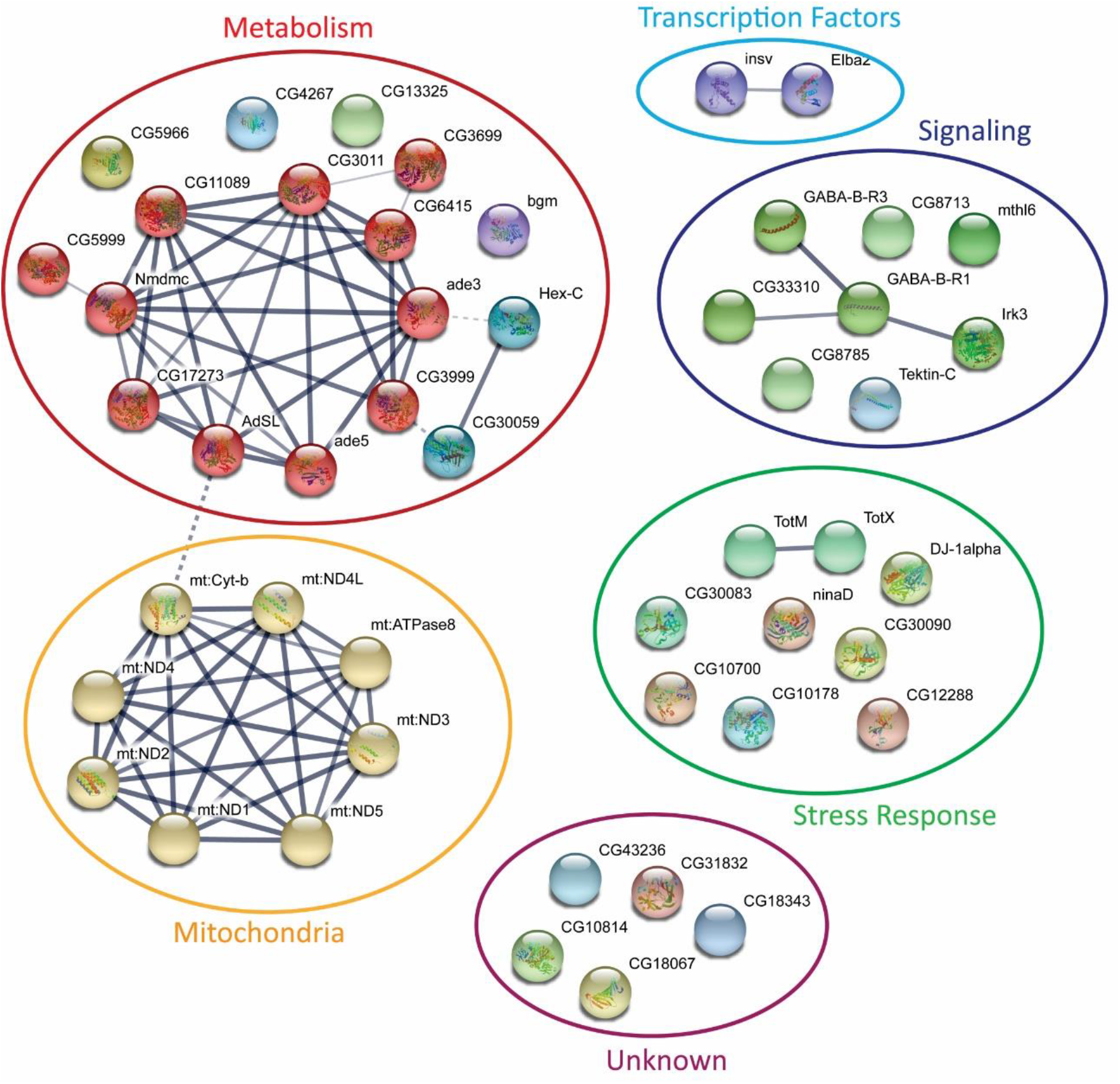
STRING interaction diagram of the genes deregulated in the *tj*-TR mutants, Related to Figure 5. Each node represents a gene that was identified to be deregulated among the three *tj*-TR mutants in high-throughput RNA sequencing analysis of brain samples. The threshold for dysregulation, upon comparing the expression levels between each mutant was set to be >+1 or <-1 log_2_ fold change. Pairwise comparisons were made between *tj^nat/nat^, tj^TR/TR^* and *tj^nTR/nTR^* in all permutations.

**Table S1.** List of candidates selected for TR validation

| Gene | TR length, codons | Region Profile | Peptide feature | Expression |
| --- | --- | --- | --- | --- |
| *br* | 131 | Ala/Gly/His rich | Disordered | Embryonic/larval CNS |
| *klu* | 15 | - | - | Embryonic neuroblasts, larval CNS |
| *chinmo* | 236 | Thr rich | BTB domain, disordered | Embryonic/larval nervous system, eye disc, adult testes |
| *wit* | 10 | - | - | Embryonic/larval/adult CNS, midgut, eye, salivary gland |
| *dsx* | 23 | - | Non-cytoplasmic/signal peptide | Embryonic gonad, embryonic/larval/adult CNS, testis |
| *Khc-73* | 58 | - | - | Enriched in larval/pupal CNS, ubiquitous |
| *fru* | 187 | Gln/Asn rich | Polar, disordered | Ubiquitous in embryos, larval/pupal/adult CNS |
| *svp* | 11 | - |  | Embryonic neuroblasts, larval photoreceptor cells, fat body, adult optic lobe, photoreceptors |
| *aPKC* | 131 | Asn/Gln rich | Polar, disordered | Ubiquitous in early embryos, larval/pupal/adult CNS |
| *dlg1* | 41 | - | - | Embryonic/larval/adult CNS, salivary glands, fat bodies |
| *tj* | 44 | - | Disordered | Gonadal somatic cells, embryonic/larval CNS |

**Table S2.** Forced TR of Tj affects germarium and GSC niche morphology

| **Genotype** | **Observed germaria phenotypes** |  |  | **p-value** | **Number of germaria analyzed** |
| --- | --- | --- | --- | --- | --- |
|  | **Normal** | **Deformed** | **Small** |  |  |
| *tj<sup>nat/nat</sup>* | 91% | 5% | 4% |  | 128 |
| *tj<sup>TR/TR</sup>* | 31% | 18% | 51% | <sup>1</sup>p=2.7e<sup>-21</sup><br><sup>2</sup>p=1.1e<sup>-15</sup> | 146 |
| *tj<sup>nTR/nTR</sup>* | 80% | 10% | 10% | <sup>1</sup>p=0.155 | 153 |
| **Genotype** | **Observed GSC niche phenotypes** |  |  | **p-value** | **Number of niches analyzed** |
|  | **Normal** | **Small& Absent** | **Large&Fused** |  |  |
| *tj<sup>nat/nat</sup>* | 96% | 2% | 2% |  | 122 |
| *tj<sup>TR/TR</sup>* | 33% | 56% | 11% | <sup>1</sup>p=8.7e<sup>-25</sup><br><sup>2</sup>p=1.5e<sup>-05</sup> | 147 |
| *tj<sup>nTR/nTR</sup>* | 88% | 9% | 3% | <sup>1</sup>p=0.215 | 138 |

For comparison of the observed germaria phenotypes, two-way tables and $\chi^2$ -test were used.
<sup>1</sup> – compared to *tj<sup>nat/nat</sup>*
<sup>2</sup> – compared to *tj<sup>nTR/nTR</sup>*

**Table S3.**
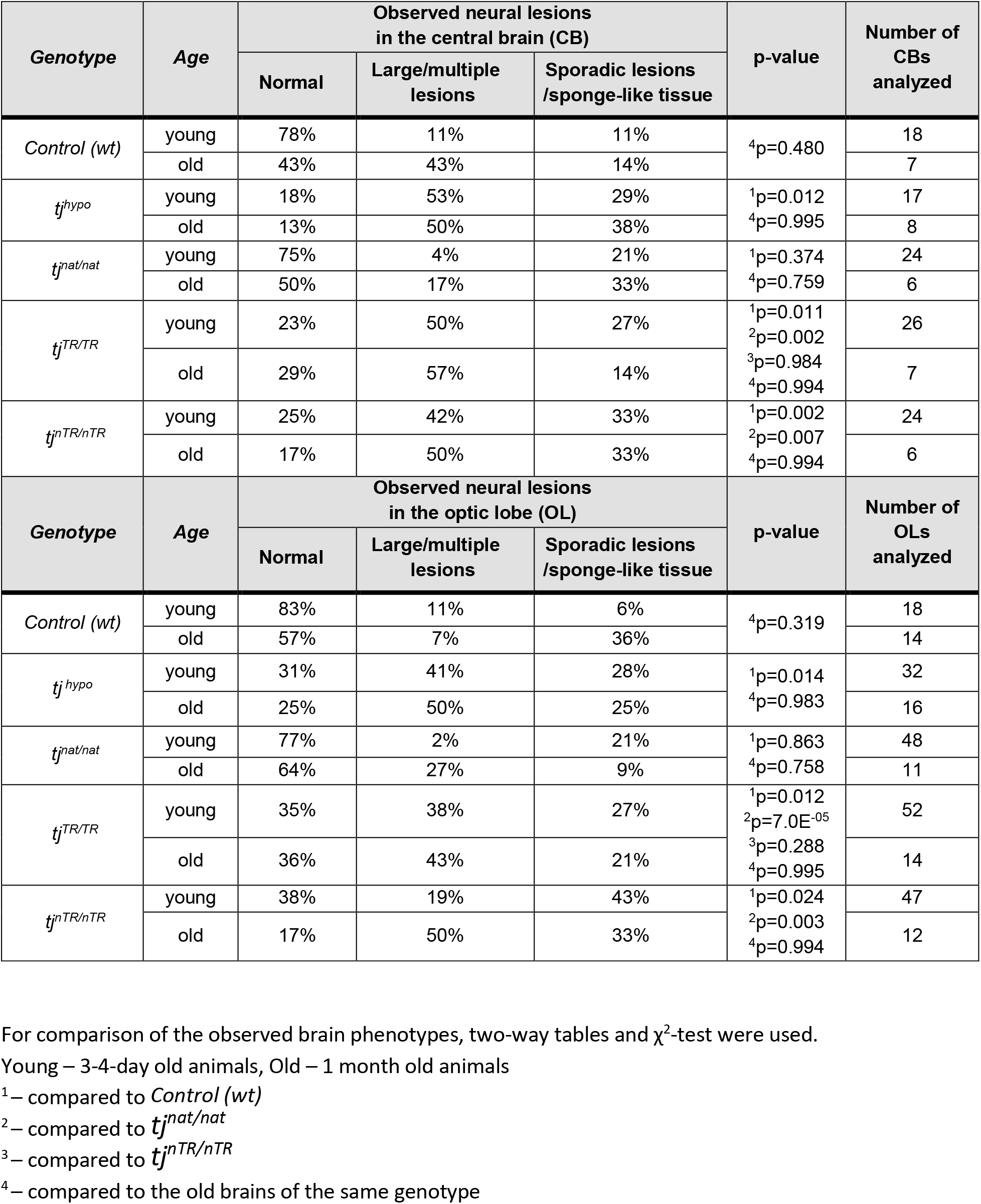
Perturbed regulation of TR of Tj causes brain neurodegeneration

**Table S4.**
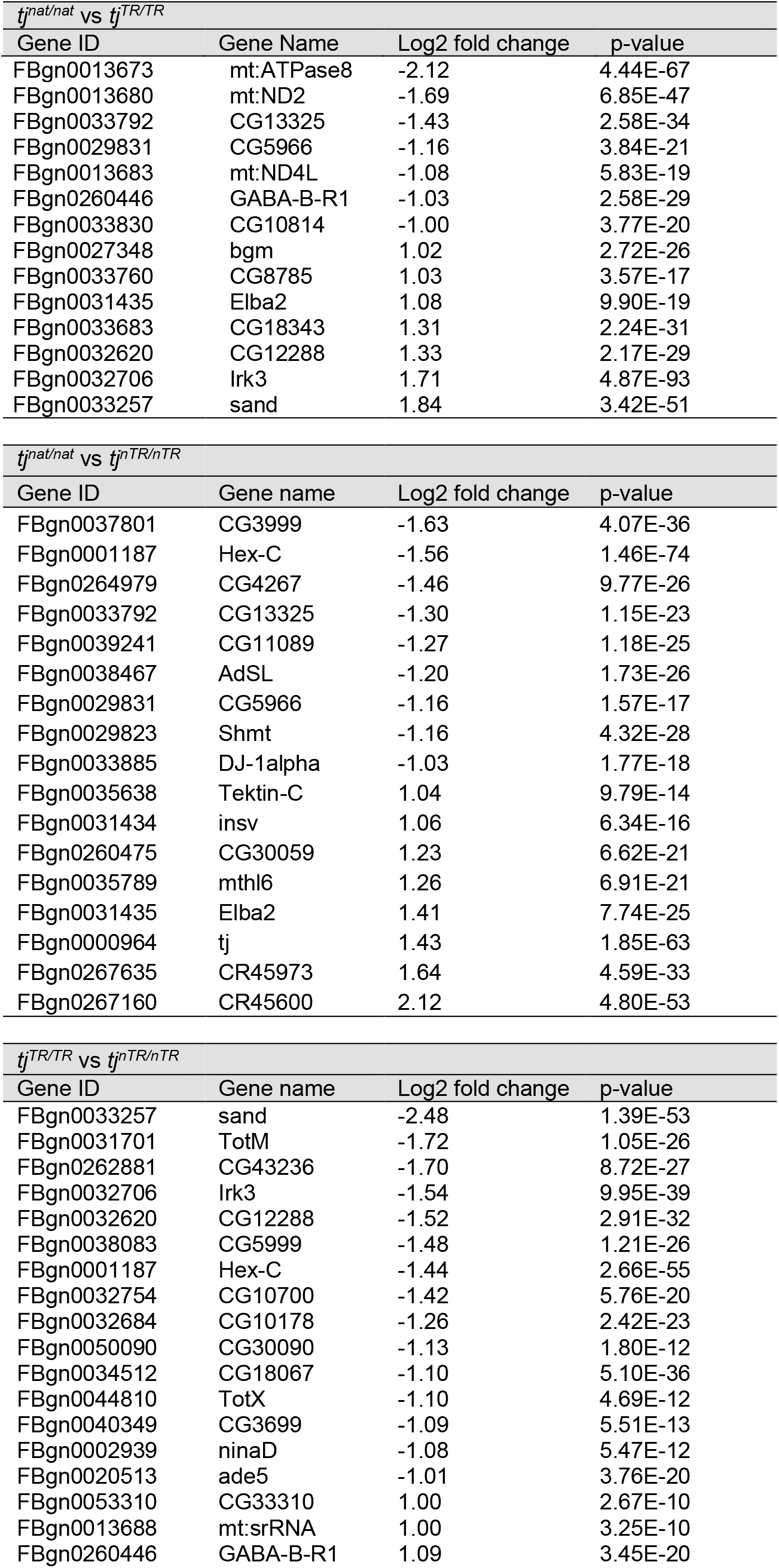
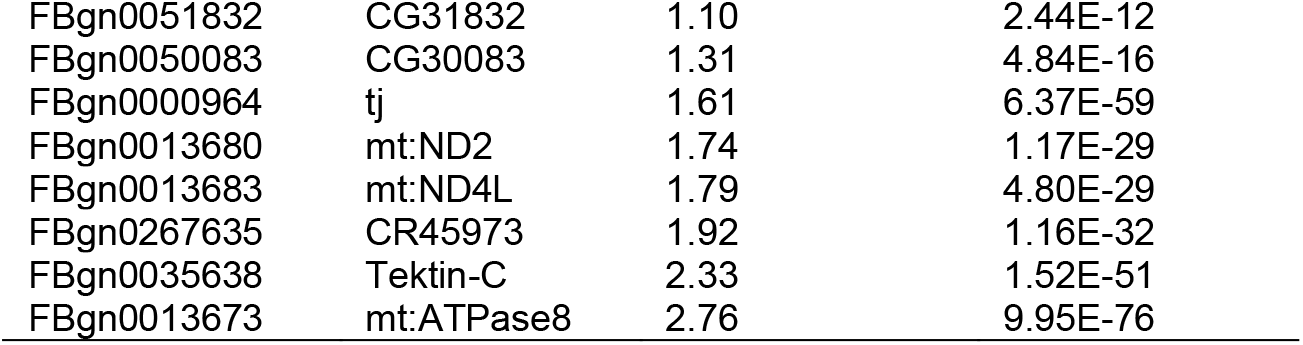
Genes identified to be dysregulated in *tj*-TR mutants

**Table S5.**
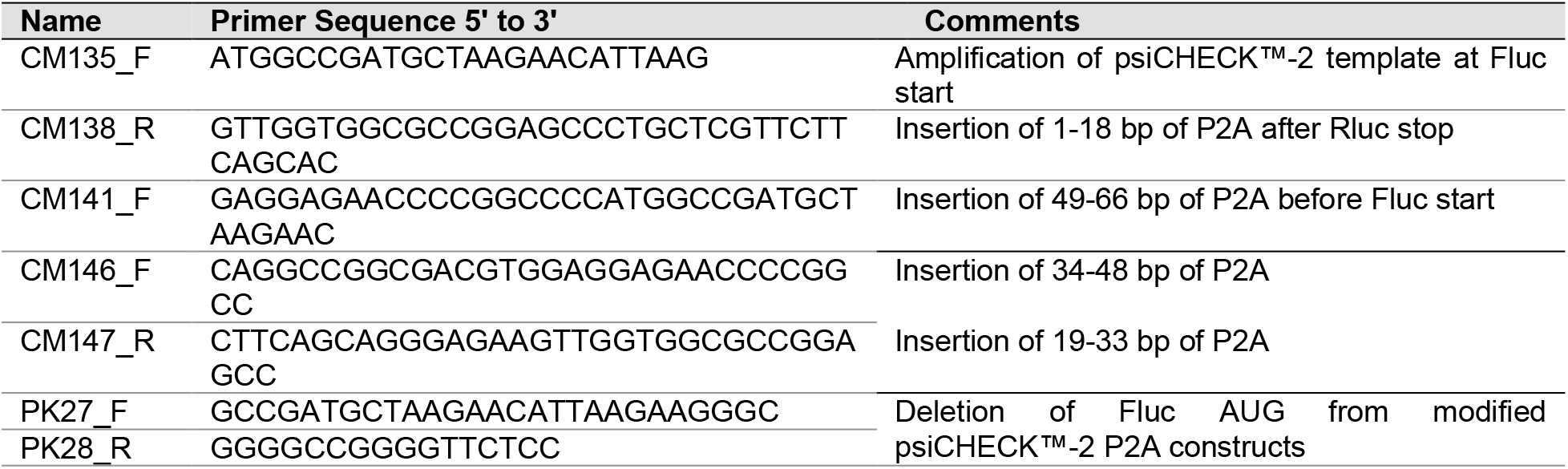
List of primers used for psiCHECK^™^-2 vector modification

**Table S6.**
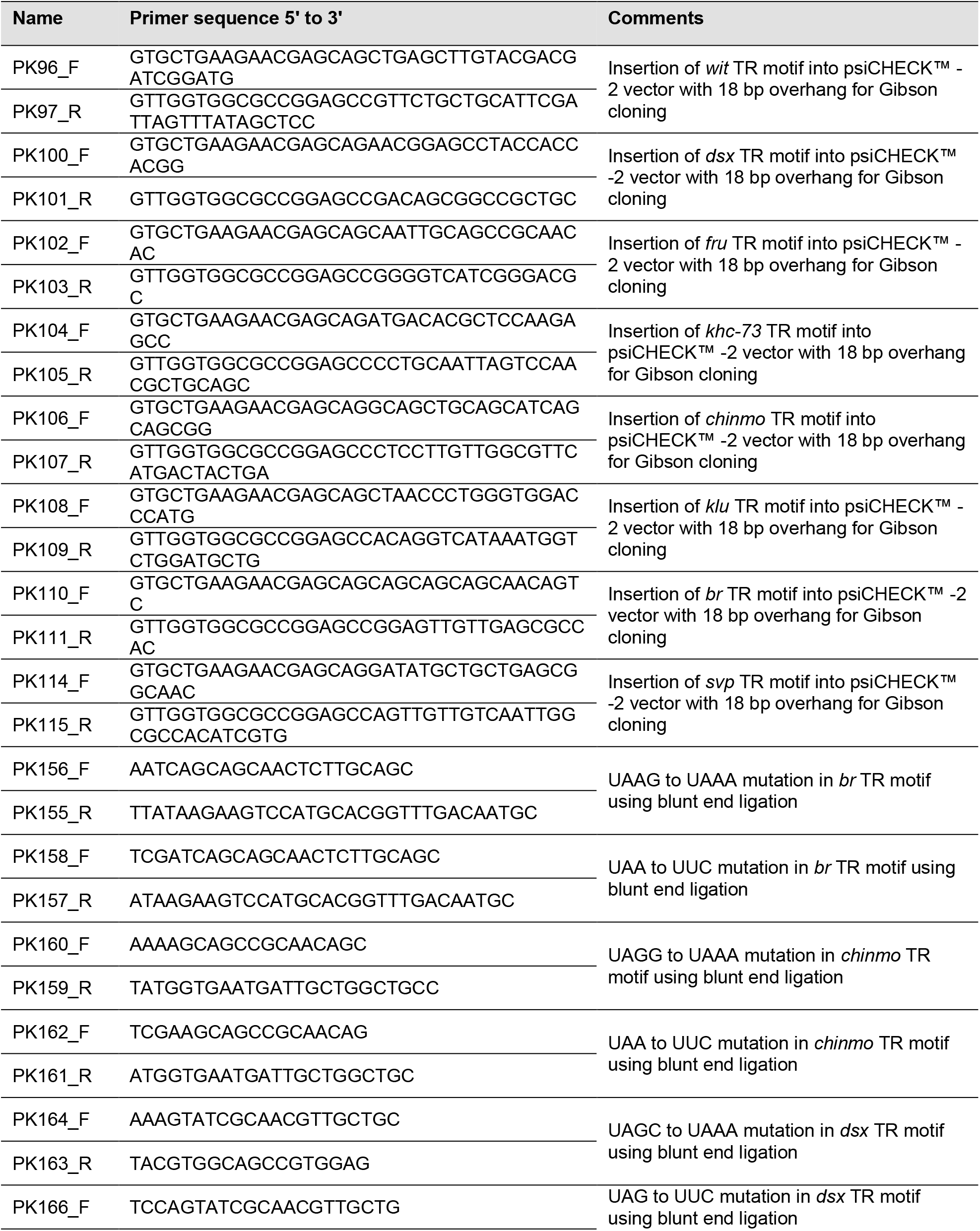
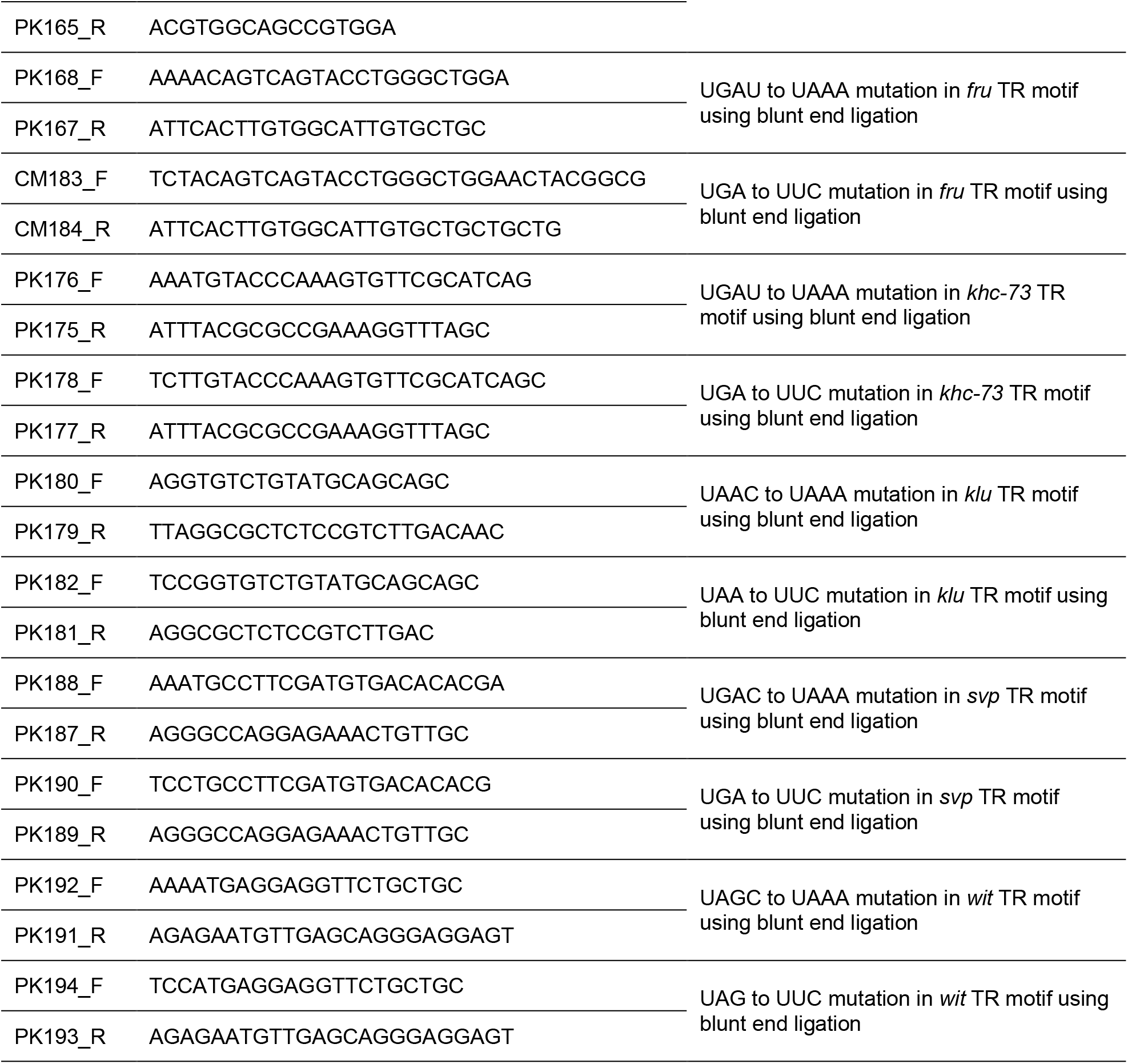
List of primers used for generating dual luciferase constructs for candidate TR genes

**Table S7.**
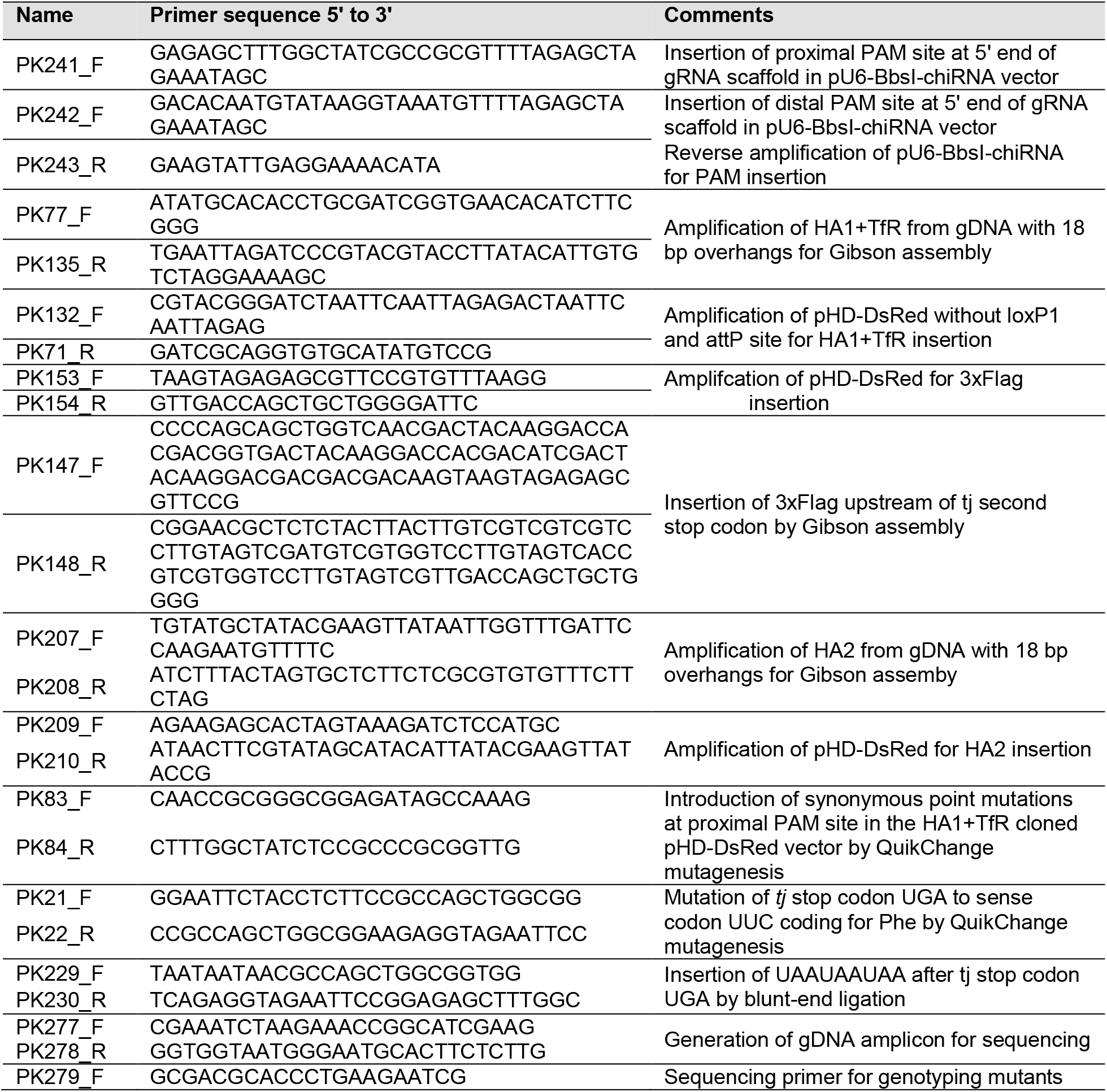
List of primers used for preparing constructs for CRISPR/Cas9 injections

**Table S8.**
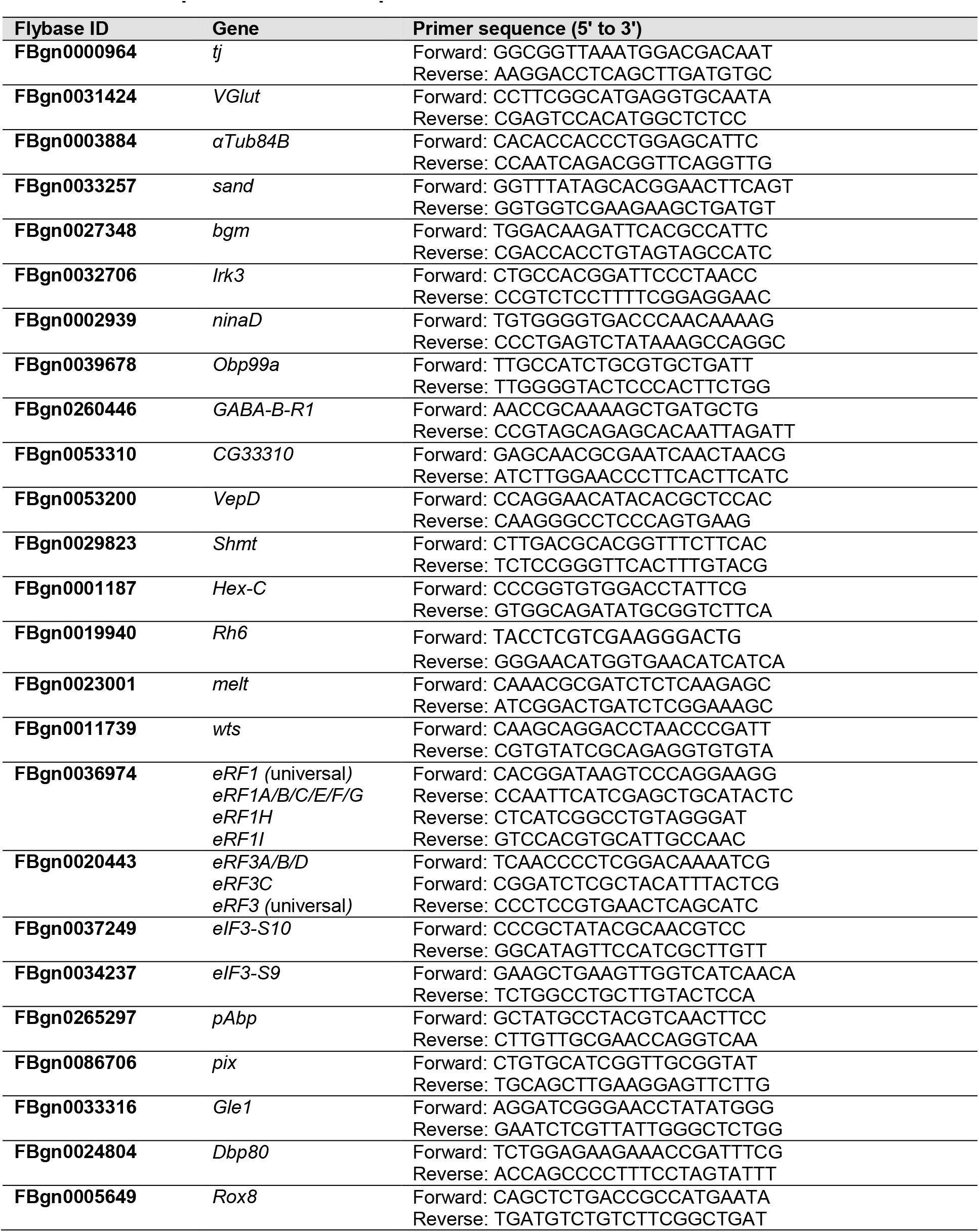
List of primers used for qPCR

**Table S9.**
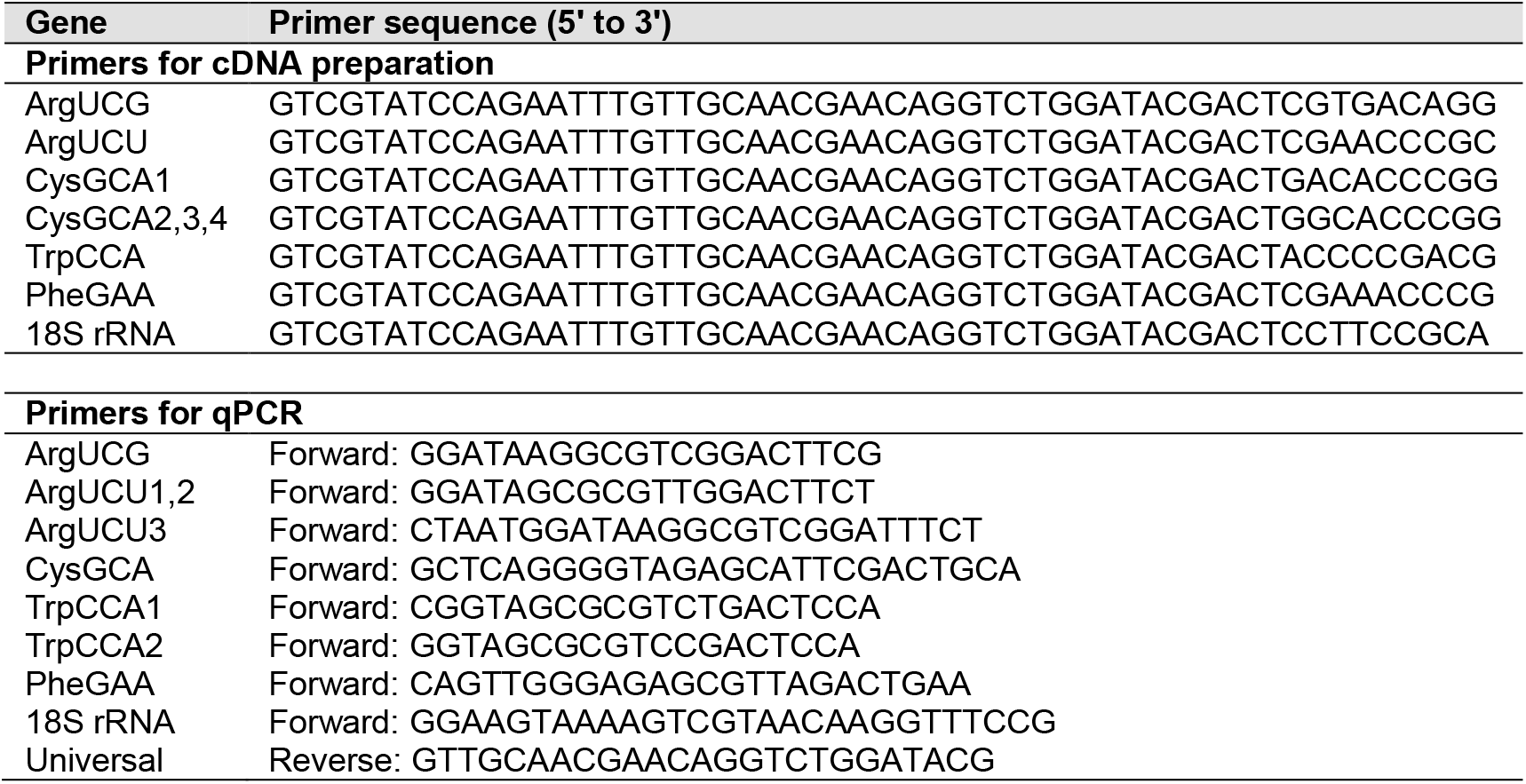
List of primers for tRNA quantification

## Notes

### Competing Interest Statement

The authors have declared no competing interest.

## References

A, P., and Weber, S.C. (2019). Evidence for and against Liquid-Liquid Phase Separation in the Nucleus. Noncoding RNA 5.

Arribere, J.A., Cenik, E.S., Jain, N., Hess, G.T., Lee, C.H., Bassik, M.C., and Fire, A.Z. (2016). Translation readthrough mitigation. Nature 534, 719–723.

Bai, J., and Montell, D. (2002). Eyes absent, a key repressor of polar cell fate during Drosophila oogenesis. Development 129, 5377–5388.

Beier, H., and Grimm, M. (2001). Misreading of termination codons in eukaryotes by natural nonsense suppressor tRNAs. Nucleic Acids Res 29, 4767–4782.

Beissel, C., Neumann, B., Uhse, S., Hampe, I., Karki, P., and Krebber, H. (2019). Translation termination depends on the sequential ribosomal entry of eRF1 and eRF3. Nucleic Acids Res 47, 4798–4813.

Bersch, K., Lobos Matthei, I., and Thoms, S. (2018). Multiple Localization by Functional Translational Readthrough. Subcell Biochem 89, 201–219.

Beznoskova, P., Gunisova, S., and Valasek, L.S. (2016). Rules of UGA-N decoding by near-cognate tRNAs and analysis of readthrough on short uORFs in yeast. Rna 22, 456–466.

Beznoskova, P., Wagner, S., Jansen, M.E., von der Haar, T., and Valasek, L.S. (2015). Translation initiation factor eIF3 promotes programmed stop codon readthrough. Nucleic Acids Res 43, 5099–5111.

Blanchet, S., Cornu, D., Argentini, M., and Namy, O. (2014). New insights into the incorporation of natural suppressor tRNAs at stop codons in Saccharomyces cerevisiae. Nucleic Acids Res 42, 10061–10072.

Bolger, T.A., Folkmann, A.W., Tran, E.J., and Wente, S.R. (2008). The mRNA export factor Gle1 and inositol hexakisphosphate regulate distinct stages of translation. Cell 134, 624–633.

Bonetti, B., Fu, L., Moon, J., and Bedwell, D.M. (1995). The efficiency of translation termination is determined by a synergistic interplay between upstream and downstream sequences in Saccharomyces cerevisiae. J Mol Biol 251, 334–345.

Carnes, J., Jacobson, M., Leinwand, L., and Yarus, M. (2003). Stop codon suppression via inhibition of eRF1 expression. RNA 9, 648–653.

Cassan, M., and Rousset, J.P. (2001). UAG readthrough in mammalian cells: effect of upstream and downstream stop codon contexts reveal different signals. BMC Mol Biol 2, 3.

Chao, A.T., Dierick, H.A., Addy, T.M., and Bejsovec, A. (2003). Mutations in eukaryotic release factors 1 and 3 act as general nonsense suppressors in Drosophila. Genetics 165, 601–612.

Chauvin, C., Salhi, S., Le Goff, C., Viranaicken, W., Diop, D., and Jean-Jean, O. (2005). Involvement of human release factors eRF3a and eRF3b in translation termination and regulation of the termination complex formation. Mol Cell Biol 25, 5801–5811.

Chen, Y., Sun, T., Bi, Z., Ni, J.Q., Pastor-Pareja, J.C., and Javid, B. (2020). Premature termination codon readthrough in Drosophila varies in a developmental and tissue-specific manner. Sci Rep 10, 8485.

Cheng, Z., Saito, K., Pisarev, A.V., Wada, M., Pisareva, V.P., Pestova, T.V., Gajda, M., Round, A., Kong, C., Lim, M., et al. (2009). Structural insights into eRF3 and stop codon recognition by eRF1. Genes Dev 23, 1106–1118.

Cridge, A.G., Crowe-McAuliffe, C., Mathew, S.F., and Tate, W.P. (2018). Eukaryotic translational termination efficiency is influenced by the 3’ nucleotides within the ribosomal mRNA channel. Nucleic Acids Res 46, 1927–1944.

Csibra, E., Brierley, I., and Irigoyen, N. (2014). Modulation of stop codon read-through efficiency and its effect on the replication of murine leukemia virus. J Virol 88, 10364–10376.

De Bellis, M., Pisani, F., Mola, M.G., Rosito, S., Simone, L., Buccoliero, C., Trojano, M., Nicchia, G.P., Svelto, M., and Frigeri, A. (2017). Translational readthrough generates new astrocyte AQP4 isoforms that modulate supramolecular clustering, glial endfeet localization, and water transport. Glia 65, 790–803.

Dunn, J.G., Foo, C.K., Belletier, N.G., Gavis, E.R., and Weissman, J.S. (2013). Ribosome profiling reveals pervasive and regulated stop codon readthrough in Drosophila melanogaster. Elife 2, e01179.

Eliazer, S., Palacios, V., Wang, Z., Kollipara, R.K., Kittler, R., and Buszczak, M. (2014). Lsd1 restricts the number of germline stem cells by regulating multiple targets in escort cells. PLoS Genet 10, e1004200.

Eswarappa, S.M., Potdar, A.A., Koch, W.J., Fan, Y., Vasu, K., Lindner, D., Willard, B., Graham, L.M., DiCorleto, P.E., and Fox, P.L. (2014). Programmed translational readthrough generates antiangiogenic VEGF-Ax. Cell 157, 1605–1618.

Fearon, K., McClendon, V., Bonetti, B., and Bedwell, D.M. (1994). Premature translation termination mutations are efficiently suppressed in a highly conserved region of yeast Ste6p, a member of the ATP-binding cassette (ABC) transporter family. J Biol Chem 269, 17802–17808.

Felsenstein, K.M., and Goff, S.P. (1988). Expression of the gag-pol fusion protein of Moloney murine leukemia virus without gag protein does not induce virion formation or proteolytic processing. J Virol 62, 2179–2182.

Feng, Y.X., Copeland, T.D., Oroszlan, S., Rein, A., and Levin, J.G. (1990). Identification of amino acids inserted during suppression of UAA and UGA termination codons at the gag-pol junction of Moloney murine leukemia virus. Proc Natl Acad Sci U S A 87, 8860–8863.

Firth, A.E., Wills, N.M., Gesteland, R.F., and Atkins, J.F. (2011). Stimulation of stop codon readthrough: frequent presence of an extended 3’ RNA structural element. Nucleic Acids Res 39, 6679–6691.

Fleming, I., and Cavalcanti, A.R.O. (2019). Selection for tandem stop codons in ciliate species with reassigned stop codons. PLoS One 14, e0225804.

Gelbart, W.M., and Emmert, D.B. (2013). FlyBase High Throughput Expression Pattern Data

Gunawan, F., Arandjelovic, M., and Godt, D. (2013). The Maf factor Traffic jam both enables and inhibits collective cell migration in Drosophila oogenesis. Development 140, 2808–2817.

Hofstetter, H., Monstein, H.J., and Weissmann, C. (1974). The readthrough protein A1 is essential for the formation of viable Q beta particles. Biochim Biophys Acta 374, 238–251.

Hoshino, S., Imai, M., Mizutani, M., Kikuchi, Y., Hanaoka, F., Ui, M., and Katada, T. (1998). Molecular cloning of a novel member of the eukaryotic polypeptide chain-releasing factors (eRF). Its identification as eRF3 interacting with eRF1. J Biol Chem 273, 22254–22259.

Howard, M.T., Shirts, B.H., Petros, L.M., Flanigan, K.M., Gesteland, R.F., and Atkins, J.F. (2000). Sequence specificity of aminoglycoside-induced stop condon readthrough: potential implications for treatment of Duchenne muscular dystrophy. Ann Neurol 48, 164–169.

Huang, A.M., Rehm, E.J., and Rubin, G.M. (2009). Quick preparation of genomic DNA from Drosophila. Cold Spring Harb Protoc 2009, pdb prot5198.

Hudson, A.M., Loughran, G., Szabo, N.L., Wills, N.M., Atkins, J.F., and Cooley, L. (2020). Tissue-specific dynamic codon redefinition in Drosophila. In bioRxiv.

Ivanov, A., Mikhailova, T., Eliseev, B., Yeramala, L., Sokolova, E., Susorov, D., Shuvalov, A., Schaffitzel, C., and Alkalaeva, E. (2016). PABP enhances release factor recruitment and stop codon recognition during translation termination. Nucleic Acids Res 44, 7766–7776.

Jakobsen, C.G., Sogaard, T.M.M., Jean-Jean, O., Frolova, L.Y., and Justesen, J. (2001). Identification of eRF3b, a human polypeptide chain release factor with eRF3 activity in vitro and in vivo. Mol Biol+ 35, 575–583.

Janzen, D.M., and Geballe, A.P. (2004). The effect of eukaryotic release factor depletion on translation termination in human cell lines. Nucleic Acids Res 32, 4491–4502.

Jukam, D., Xie, B., Rister, J., Terrell, D., Charlton-Perkins, M., Pistillo, D., Gebelein, B., Desplan, C., and Cook, T. (2013). Opposite feedbacks in the Hippo pathway for growth control and neural fate. Science 342, 1238016.

Jungreis, I., Chan, C.S., Waterhouse, R.M., Fields, G., Lin, M.F., and Kellis, M. (2016). Evolutionary Dynamics of Abundant Stop Codon Readthrough. Mol Biol Evol 33, 3108–3132.

Jungreis, I., Lin, M.F., Spokony, R., Chan, C.S., Negre, N., Victorsen, A., White, K.P., and Kellis, M. (2011). Evidence of abundant stop codon readthrough in Drosophila and other metazoa. Genome Res 21, 2096–2113.

Kataoka, K., Noda, M., and Nishizawa, M. (1994). Maf nuclear oncoprotein recognizes sequences related to an AP-1 site and forms heterodimers with both Fos and Jun. Mol Cell Biol 14, 700–712.

Keeling, K.M., Lanier, J., Du, M., Salas-Marco, J., Gao, L., Kaenjak-Angeletti, A., and Bedwell, D.M. (2004). Leaky termination at premature stop codons antagonizes nonsense-mediated mRNA decay in S. cerevisiae. RNA 10, 691–703.

Kim, E., Magen, A., and Ast, G. (2007). Different levels of alternative splicing among eukaryotes. Nucleic Acids Res 35, 125–131.

Klagges, B.R., Heimbeck, G., Godenschwege, T.A., Hofbauer, A., Pflugfelder, G.O., Reifegerste, R., Reisch, D., Schaupp, M., Buchner, S., and Buchner, E. (1996). Invertebrate synapsins: a single gene codes for several isoforms in Drosophila. J Neurosci 16, 3154–3165.

Kleppe, A.S., and Bornberg-Bauer, E. (2018). Robustness by intrinsically disordered C-termini and translational readthrough. Nucleic Acids Res 46, 10184–10194.

Konig, A., and Shcherbata, H.R. (2015). Soma influences GSC progeny differentiation via the cell adhesion-mediated steroid-let-7-Wingless signaling cascade that regulates chromatin dynamics. Biol Open 4, 285–300.

Kononenko, A.V., Mitkevich, V.A., Dubovaya, V.I., Kolosov, P.M., Makarov, A.A., and Kisselev, L.L. (2008). Role of the individual domains of translation termination factor eRF1 in GTP binding to eRF3. Proteins 70, 388–393.

Konstantinides, N., Kapuralin, K., Fadil, C., Barboza, L., Satija, R., and Desplan, C. (2018). Phenotypic Convergence: Distinct Transcription Factors Regulate Common Terminal Features. Cell 174, 622–635 e613.

Kornblihtt, A.R., Schor, I.E., Allo, M., Dujardin, G., Petrillo, E., and Munoz, M.J. (2013). Alternative splicing: a pivotal step between eukaryotic transcription and translation. Nat Rev Mol Cell Biol 14, 153–165.

Korniy, N., Goyal, A., Hoffmann, M., Samatova, E., Peske, F., Pohlmann, S., and Rodnina, M.V. (2019). Modulation of HIV-1 Gag/Gag-Pol frameshifting by tRNA abundance. Nucleic Acids Res 47, 5210–5222.

Kucherenko, M.M., Marrone, A.K., Rishko, V.M., Yatsenko, A.S., Klepzig, A., and Shcherbata, H.R. (2010). Paraffin-embedded and frozen sections of Drosophila adult muscles. J Vis Exp.

Kurokawa, H., Motohashi, H., Sueno, S., Kimura, M., Takagawa, H., Kanno, Y., Yamamoto, M., and Tanaka, T. (2009). Structural basis of alternative DNA recognition by Maf transcription factors. Mol Cell Biol 29, 6232–6244.

Lai, C.M., Lin, K.Y., Kao, S.H., Chen, Y.N., Huang, F., and Hsu, H.J. (2017). Hedgehog signaling establishes precursors for germline stem cell niches by regulating cell adhesion. J Cell Biol 216, 1439–1453.

Li, M., Hu, X., Zhang, S., Ho, M.S., Wu, G., and Zhang, L. (2019). Traffic jam regulates the function of the ovarian germline stem cell progeny differentiation niche during pre-adult stage in Drosophila. Sci Rep 9, 10124.

Li, M.A., Alls, J.D., Avancini, R.M., Koo, K., and Godt, D. (2003). The large Maf factor Traffic Jam controls gonad morphogenesis in Drosophila. Nat Cell Biol 5, 994–1000.

Loughran, G., Chou, M.Y., Ivanov, I.P., Jungreis, I., Kellis, M., Kiran, A.M., Baranov, P.V., and Atkins, J.F. (2014). Evidence of efficient stop codon readthrough in four mammalian genes. Nucleic Acids Res 42, 8928–8938.

Manuvakhova, M., Keeling, K., and Bedwell, D.M. (2000). Aminoglycoside antibiotics mediate contextdependent suppression of termination codons in a mammalian translation system. RNA 6, 1044–1055.

McCaughan, K.K., Brown, C.M., Dalphin, M.E., Berry, M.J., and Tate, W.P. (1995). Translational termination efficiency in mammals is influenced by the base following the stop codon. Proc Natl Acad Sci U S A 92, 5431–5435.

Mikhailova, T., Shuvalova, E., Ivanov, A., Susorov, D., Shuvalov, A., Kolosov, P.M., and Alkalaeva, E. (2017). RNA helicase DDX19 stabilizes ribosomal elongation and termination complexes. Nucleic Acids Res 45, 1307–1318.

Namy, O., Duchateau-Nguyen, G., Hatin, I., Hermann-Le Denmat, S., Termier, M., and Rousset, J.P. (2003). Identification of stop codon readthrough genes in Saccharomyces cerevisiae. Nucleic Acids Res 31, 2289–2296.

Namy, O., Duchateau-Nguyen, G., and Rousset, J.P. (2002). Translational readthrough of the PDE2 stop codon modulates cAMP levels in Saccharomyces cerevisiae. Mol Microbiol 43, 641–652.

Napthine, S., Yek, C., Powell, M.L., Brown, T.D., and Brierley, I. (2012). Characterization of the stop codon readthrough signal of Colorado tick fever virus segment 9 RNA. RNA 18, 241–252.

Owen, I., and Shewmaker, F. (2019). The Role of Post-Translational Modifications in the Phase Transitions of Intrinsically Disordered Proteins. Int J Mol Sci 20.

Palazzo, C., Abbrescia, P., Valente, O., Nicchia, G.P., Banitalebi, S., Amiry-Moghaddam, M., Trojano, M., and Frigeri, A. (2020). Tissue Distribution of the Readthrough Isoform of AQP4 Reveals a Dual Role of AQP4ex Limited to CNS. Int J Mol Sci 21.

Panchal, T., Chen, X., Alchits, E., Oh, Y., Poon, J., Kouptsova, J., Laski, F.A., and Godt, D. (2017). Specification and spatial arrangement of cells in the germline stem cell niche of the Drosophila ovary depend on the Maf transcription factor Traffic jam. PLoS Genet 13, e1006790.

Pancsa, R., Macossay-Castillo, M., Kosol, S., and Tompa, P. (2016). Computational analysis of translational readthrough proteins in Drosophila and yeast reveals parallels to alternative splicing. Sci Rep 6, 32142.

Pelham, H.R. (1978). Leaky UAG termination codon in tobacco mosaic virus RNA. Nature 272, 469–471.

Preis, A., Heuer, A., Barrio-Garcia, C., Hauser, A., Eyler, D.E., Berninghausen, O., Green, R., Becker, T., and Beckmann, R. (2014). Cryoelectron microscopic structures of eukaryotic translation termination complexes containing eRF1-eRF3 or eRF1-ABCE1. Cell Rep 8, 59–65.

Raj, B., and Blencowe, B.J. (2015). Alternative Splicing in the Mammalian Nervous System: Recent Insights into Mechanisms and Functional Roles. Neuron 87, 14–27.

Robinson, D.N., and Cooley, L. (1997). Examination of the function of two kelch proteins generated by stop codon suppression. Development 124, 1405–1417.

Roy, B., Leszyk, J.D., Mangus, D.A., and Jacobson, A. (2015). Nonsense suppression by near-cognate tRNAs employs alternative base pairing at codon positions 1 and 3. Proc Natl Acad Sci U S A 112, 3038–3043.

Saito, K., Inagaki, S., Mituyama, T., Kawamura, Y., Ono, Y., Sakota, E., Kotani, H., Asai, K., Siomi, H., and Siomi, M.C. (2009). A regulatory circuit for piwi by the large Maf gene traffic jam in Drosophila. Nature 461, 1296–1299.

Samson, M.L., Lisbin, M.J., and White, K. (1995). Two distinct temperature-sensitive alleles at the elav locus of Drosophila are suppressed nonsense mutations of the same tryptophan codon. Genetics 141, 1101–1111.

Sapkota, D., Lake, A.M., Yang, W., Yang, C., Wesseling, H., Guise, A., Uncu, C., Dalal, J.S., Kraft, A.W., Lee, J.M., et al. (2019). Cell-Type-Specific Profiling of Alternative Translation Identifies Regulated Protein Isoform Variation in the Mouse Brain. Cell Rep 26, 594–607 e597.

Shcherbata, H.R., Yatsenko, A.S., Patterson, L., Sood, V.D., Nudel, U., Yaffe, D., Baker, D., and Ruohola-Baker, H. (2007). Dissecting muscle and neuronal disorders in a Drosophila model of muscular dystrophy. EMBO J 26, 481–493.

Shoemaker, C.J., and Green, R. (2011). Kinetic analysis reveals the ordered coupling of translation termination and ribosome recycling in yeast. Proc Natl Acad Sci U S A 108, E1392–1398.

Steneberg, P., and Samakovlis, C. (2001). A novel stop codon readthrough mechanism produces functional Headcase protein in Drosophila trachea. EMBO Rep 2, 593–597.

Su, C.H., D, D., and Tarn, W.Y. (2018). Alternative Splicing in Neurogenesis and Brain Development. Front Mol Biosci 5, 12.

Tian, B., and Manley, J.L. (2017). Alternative polyadenylation of mRNA precursors. Nat Rev Mol Cell Biol 18, 18–30.

Touriol, C., Bornes, S., Bonnal, S., Audigier, S., Prats, H., Prats, A.C., and Vagner, S. (2003). Generation of protein isoform diversity by alternative initiation of translation at non-AUG codons. Biol Cell 95, 169–178.

Trotta, E. (2016). Selective forces and mutational biases drive stop codon usage in the human genome: a comparison with sense codon usage. BMC Genomics 17, 366.

Urakov, V.N., Mitkevich, O.V., Safenkova, I.V., and Ter-Avanesyan, M.D. (2017). Ribosome-bound Pub1 modulates stop codon decoding during translation termination in yeast. FEBS J 284, 1914–1930.

Urban, C., and Beier, H. (1995). Cysteine tRNAs of plant origin as novel UGA suppressors. Nucleic Acids Res 23, 4591–4597.

Urban, C., Zerfass, K., Fingerhut, C., and Beier, H. (1996). UGA suppression by tRNACmCATrp occurs in diverse virus RNAs due to a limited influence of the codon context. Nucleic Acids Res 24, 3424–3430.

Venables, J.P., Tazi, J., and Juge, F. (2012). Regulated functional alternative splicing in Drosophila. Nucleic Acids Res 40, 1–10.

Wan Makhtar, W.R., Browne, G., Karountzos, A., Stevens, C., Alghamdi, Y., Bottrill, A.R., Mistry, S., Smith, E., Bushel, M., Pringle, J.H., et al. (2017). Short stretches of rare codons regulate translation of the transcription factor ZEB2 in cancer cells. Oncogene 36, 6640–6648.

Williams, I., Richardson, J., Starkey, A., and Stansfield, I. (2004). Genome-wide prediction of stop codon readthrough during translation in the yeast Saccharomyces cerevisiae. Nucleic Acids Res 32, 6605–6616.

Wingert, L., and DiNardo, S. (2015). Traffic jam functions in a branched pathway from Notch activation to niche cell fate. Development 142, 2268–2277.

Xue, F., and Cooley, L. (1993). kelch encodes a component of intercellular bridges in Drosophila egg chambers. Cell 72, 681–693.

Yatsenko, A.S., and Shcherbata, H.R. (2018). Stereotypical architecture of the stem cell niche is spatiotemporally established by miR-125-dependent coordination of Notch and steroid signaling. Development 145.

Zerfass, K., and Beier, H. (1992). The leaky UGA termination codon of tobacco rattle virus RNA is suppressed by tobacco chloroplast and cytoplasmic tRNAs(Trp) with CmCA anticodon. EMBO J 11, 4167–4173.

